# Transcriptional mediators of treatment resistance in lethal prostate cancer

**DOI:** 10.1101/2020.03.19.998450

**Authors:** Meng Xiao He, Michael S. Cuoco, Jett Crowdis, Alice Bosma-Moody, Zhenwei Zhang, Kevin Bi, Abhay Kanodia, Mei-Ju Su, Christopher Rodman, Laura DelloStritto, Parin Shah, Kelly P. Burke, Benjamin Izar, Ziad Bakouny, Alok K. Tewari, David Liu, Sabrina Y. Camp, Natalie I. Vokes, Jihye Park, Sébastien Vigneau, Lawrence Fong, Orit Rozenblatt-Rosen, Aviv Regev, Asaf Rotem, Mary-Ellen Taplin, Eliezer M. Van Allen

## Abstract

Metastatic castration resistant prostate cancer (mCRPC) is primarily treated with therapies that prevent transcriptional activity of the androgen receptor (AR), cause DNA damage, or prevent cell division. Clinical resistance to these therapies, including second-generation androgen-targeting compounds such as enzalutamide and abiraterone, is nearly universal. Other treatment modalities, including immune checkpoint inhibitors, have provided minimal benefit except in rare subsets of patients^1,2^. Both tumour intrinsic and extrinsic cellular programs contributing to therapeutic resistance remain areas of active investigation. Here we use full-length single-cell RNA-sequencing (scRNA-seq) to identify the transcriptional states of cancer and immune cells in the mCRPC microenvironment. Within cancer cells, we identified transcriptional patterns that mediate a significant proportion of inherited risk for prostate cancer, extensive heterogeneity in *AR* splicing within and between tumours, and vastly divergent regulatory programs between adenocarcinoma and small cell carcinoma. Moreover, upregulation of TGF-β signalling and epithelial-mesenchymal transition (EMT) were both associated with resistance to enzalutamide. We found that some lymph node metastases, but no bone metastases, were heavily infiltrated by dysfunctional CD8^+^ T cells, including cells undergoing dramatic clonal expansion during enzalutamide treatment. Our findings suggest avenues for rational therapeutic approaches targeting both tumour-intrinsic and immunological pathways to combat resistance to current treatment options.

---

Despite advances in targeting androgen receptor signalling and other drivers, mCRPC is typically lethal^2^. The identities and proportions of cells within human mCRPC niches is largely unknown. By defining treatment resistant states in human mCRPC, we may reveal biological drivers that inform new treatment strategies. Thus, we collected fresh biopsies from mCRPC patients from representative metastatic sites for whole exome sequencing, bulk RNA-seq, and scRNA-seq using the Smart-seq2 protocol, which generates full-length transcript sequences^3^. At time of biopsy, patients had experienced varied treatment histories, with approximately even representation before and after treatment with enzalutamide. Smaller proportions of patients had experienced abiraterone, taxanes, and other therapies (Fig. 1a). In addition to adenocarcinomas, one biopsied tumour (09171135) had a small cell carcinoma histology.

**Figure 1.**
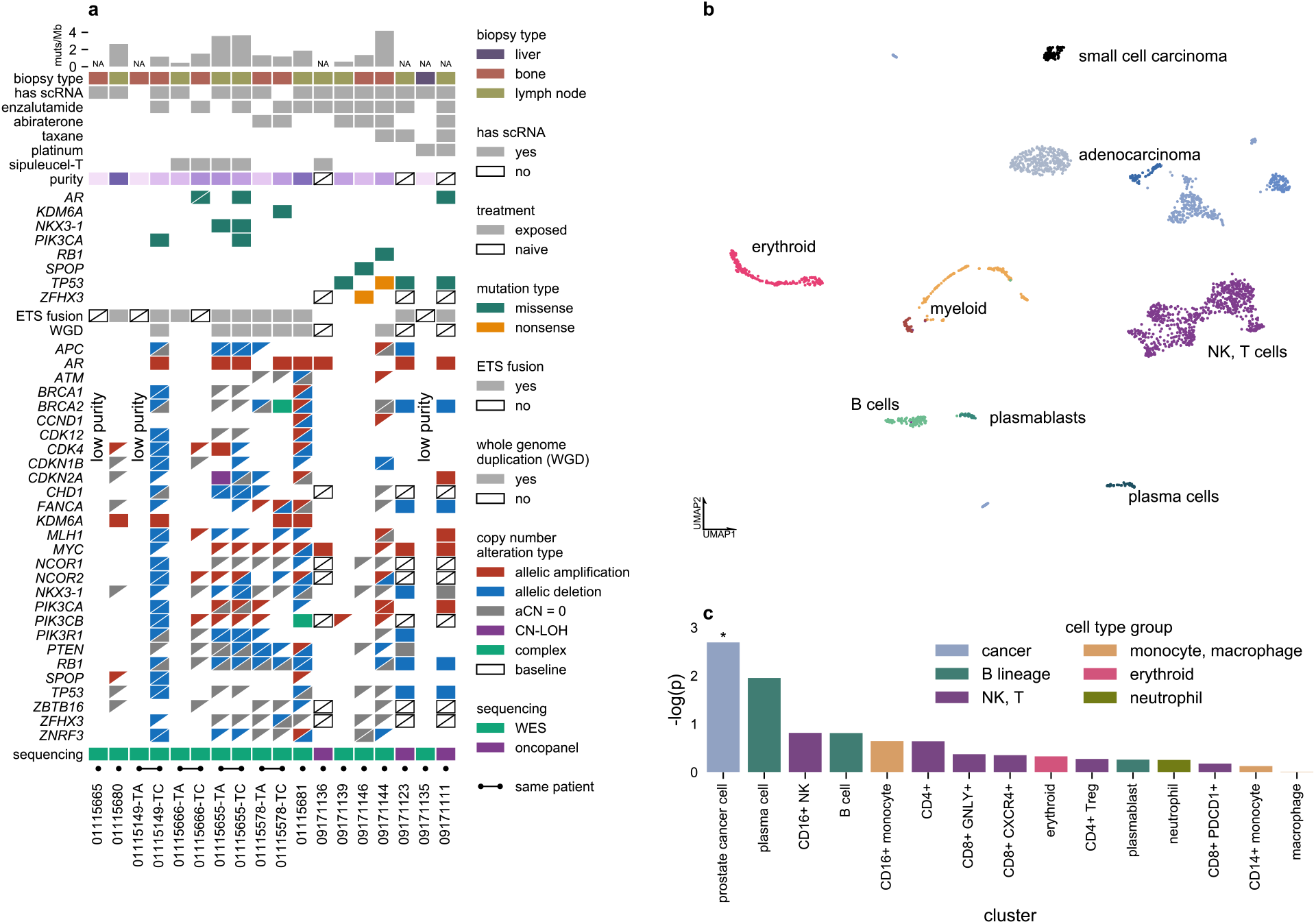
Cellular atlas of mCRPC, identifying heritability for prostate cancer enriched near genes specifically expressed in prostate cancer cells. **a)** Summary of clinical and select genomics features of patients and biopsies forming the study cohort. Each column represents a single biopsy. Where available, multiple biopsies from the same patient are displayed in adjacent columns. Patients are identified by numerical prefix, while suffixes after a dash, when present, identify biopsies from the same patient. Boxes with diagonal slashes indicate missing data, e.g. for genes not included in OncoPanel. **b)** Projection of single-cell expression onto the first two dimensions of UMAP space. Each dot represents a single cell, and colours correspond to clusters identified by the Louvain algorithm. Clusters are manually labelled with dominant cell type(s) inferred from cluster-specific expression of marker genes. **c)** Enrichment of heritability for prostate cancer near genes specifically expressed in each cell type (compared to cell types in other cell type groups). *: Benjamini-Hochberg FDR < 0.05

After quality control, our cohort consisted of 2,170 deeply sequenced cells from 14 patients and 15 biopsies, including cells from both before and after enzalutamide treatment for one patient (01115655) (Methods; Supplementary Fig. 1a). Following clustering of the single-cell transcriptomes, we manually labelled cell clusters for dominant cell type based on cluster-specific expression of marker genes (Fig. 1b; Methods; Supplementary Table 2). Cancer cells, represented in multiple clusters marked by expression of the adenocarcinoma markers *AR* and *KLK3* (which encodes prostate-specific antigen) or the neuroendocrine marker *CHGA,* were recovered from 12 biopsies, comprising over a third of the cells (n=836). The remainder included cells from the B cell lineage, natural killer (NK) and T cells, monocytes and macrophages, erythroid cells, and neutrophils.

Prostate cancer is highly heritable, with an estimated 57% of variation in risk attributed to inherited variants^4^. Genome wide association studies (GWAS) have not only identified significant risk alleles but also generated results that allow the analysis of even non-significantly associated variants in aggregate to link risk to subsets of the genome. We sought to identify cell types relevant to prostate cancer development by integrating cell-type specific expression patterns from our scRNA-seq data with results from a recent large-scale GWAS of prostate cancer risk^5^. Using LD score regression applied to specifically expressed genes (LDSC-SEG), we identified significant enrichment of germline heritability for prostate cancer in genomic intervals near genes that were specifically expressed in cancer cells (*q* = 0.031, Benjamini-Hochberg) (Fig. 1c; Methods)^6^. No significant enrichment was observed for any other cell type, indicating that when assessed during advanced disease, inherited risk for prostate cancer is primarily mediated through tumour-intrinsic mechanisms.

## Complex androgen receptor splicing

We therefore assessed transcriptional programs in cancer cells across metastatic niches and clinical contexts. As prostate adenocarcinomas are dependent on androgen signalling for survival, significant attention has been focused on the description and detection of a diverse set of *AR* splice variants. The AR protein contains a DNA-binding domain with transcriptional regulatory activity and a ligand-binding domain required for control of its activity by androgens. Splice variants that omit the ligand-binding domain, particularly AR-V7, have been hypothesized to constitutively activate downstream transcriptional programs independent of androgen binding, providing a resistance mechanism to second generation androgen-targeting therapies^7,8^. Taking advantage of our dataset’s even sequencing coverage along transcripts, we detected the presence of specific *AR* splice variants. First, we curated a transcriptome annotation of literature described isoforms (Methods). Then, we remapped all reads from individual cancer cells initially mapping to the *AR* locus, counting the number of reads that uniquely map to individual isoforms (Fig. 2a; Methods). We detected isoform-informative reads indicating the presence of many previously described splice variants within our clinical biopsies, with AR-45, AR-V7, and AR-V12 being uniquely identified in the most cells. AR-45 was detected in every biopsy with any isoform-specific reads. AR-V7 was present in biopsies from both before and after enzalutamide exposure. Strikingly, we detected multiple *AR* splice variants within the same biopsy and even within the same cell, highlighting the complexity of *AR* splicing in mCRPC.

**Figure 2.**
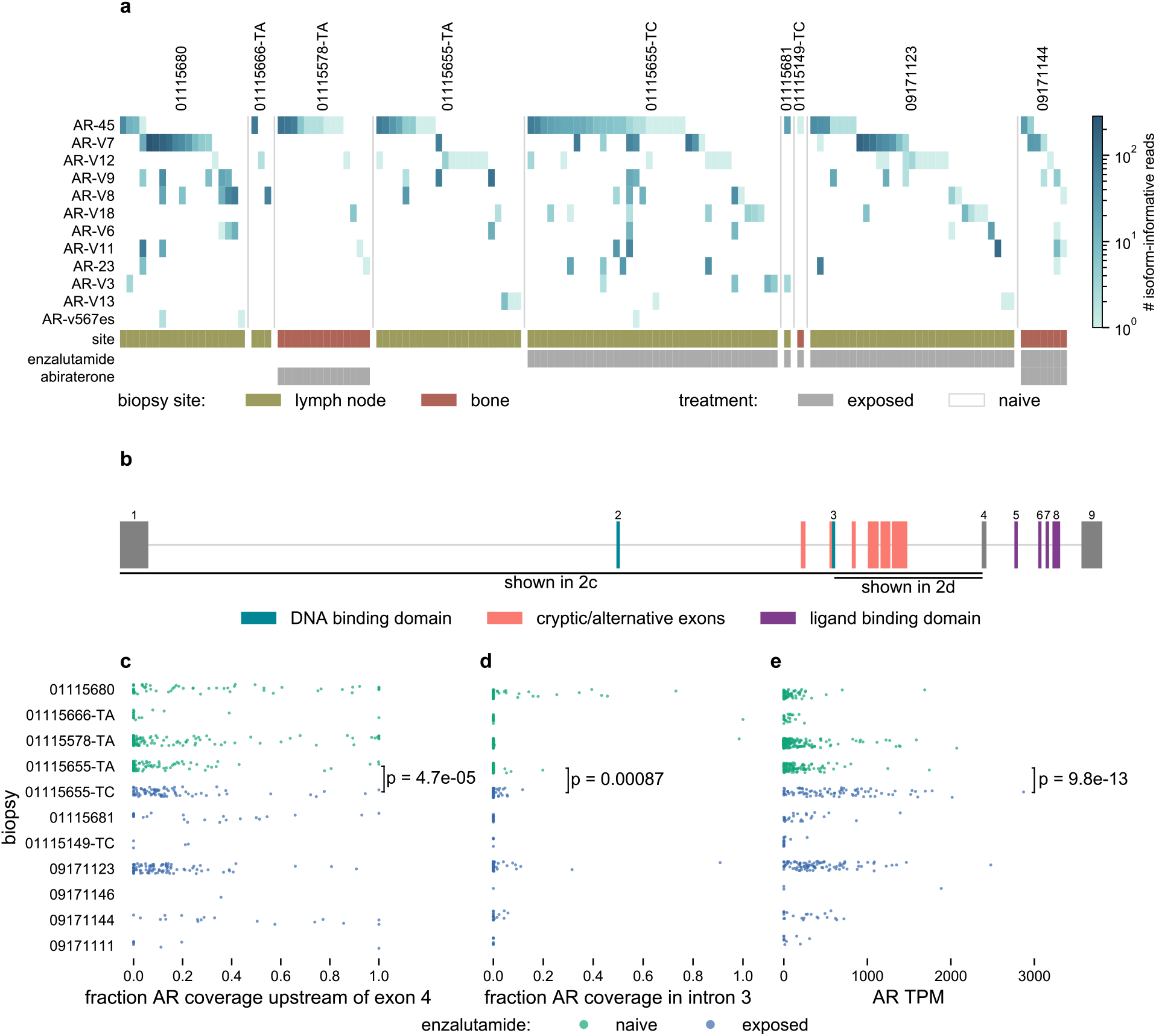
*AR* splicing varies widely across cells within the same tumour and across treatment resistance states. **a)** Heatmap displaying number of isoform-informative reads mapping to *AR* variants from single cells. Each column represents *AR* variants detected in a single cell, with only cells that had at least one isoform-informative read shown. **b)** Schematic representation of *AR* locus. Rectangles indicate exons. Exons corresponding to the full-length *AR* transcript are numbered, with exons comprising different functional domains coloured. Select alternative exons included in *AR* splice variants are indicated. **c)** Fraction of total *AR* coverage upstream of exon 4 (including the DNA-binding domain but excluding the ligand-binding domain) in single cells. **d)** Fraction of total *AR* coverage in intron 3 (including multiple cryptic/alternative exons included in truncated splice variants) in single cells. **e)** Total *AR* expression in single cells. **c, d, e)** *P* value compares cells before (n = 112) and after (n = 83) enzalutamide treatment for patient 01115655 (two-sided Mann-Whitney *U* test).

Isoform-informative reads comprise only a small fraction of reads mapping to any gene, and *AR* splice variants described in literature may not represent a complete census of all isoforms expressed *in vivo* (Supplementary Fig. 2). Therefore, we defined two alternative summary measures of *AR* splicing that permitted characterization within more of the individual cancer cells. *AR* intron 3 contains many of the terminal cryptic/alternative exons included in truncated *AR* isoforms lacking the ligand-binding domain, including AR-V7^7^. We quantified the proportion of total *AR* coverage that lies in intron 3 or in a larger interval that includes intron 3 and upstream exons, which encode the DNA-binding domain (Fig. 2b). Again, we detected significant variation between cancer cells within the same biopsy. Moreover, we detected a clear increase in both measures after enzalutamide treatment for patient 01115655, suggesting decreased transcription of full-length *AR* compared to truncating variants after treatment (Fig. 2c,d). Overall, *AR* splicing patterns in mCRPC cells were highly heterogeneous between and within tumours regardless of treatment resistance state.

## Enzalutamide resistance programs

Resistance to second generation androgen-targeting therapies poses a major clinical challenge, and previous work based on bulk whole exome and transcriptome sequencing have identified alterations in *RB1, TP53*, and *AR* as associated with poor outcomes^9^. Taking advantage of the single-cell resolution of our data, we examined cancer cells in our cohort to identify changes in expression in cells naïve and exposed to enzalutamide, which functions as a competitive inhibitor of AR that prevents nuclear localization and downstream transcriptional regulatory activity within cancer cells^10^. We scored cancer cells for expression of the MSigDB hallmark gene sets and select literature-derived gene sets, including several reported as mediating resistance mechanisms, such as genes regulated by the glucocorticoid receptor or AR-V7 and genes associated with a neuroendocrine phenotype^11–22^ (Methods). Compared to enzalutamide-naïve cells, exposed cells upregulated several MSigDB hallmark gene sets, including for EMT and TGF-β signalling (Fig. 3a,b; Supplementary Table 1). We sought to corroborate these findings in a published cohort of bulk-sequenced mCRPC transcriptomes and found a similar effect for TGF-β signalling upregulation in enzalutamide-exposed lymph node biopsies, although the number of exposed biopsies was small, and the effect was not statistically significant (Fig. 3c)^9^. We could not analyse bone biopsies due to scarcity of post-enzalutamide samples, and EMT scores were confounded with tumour purity, limiting our ability to draw conclusions from bulk sequencing for this specific finding (Supplementary Fig. 3).

**Figure 3.**
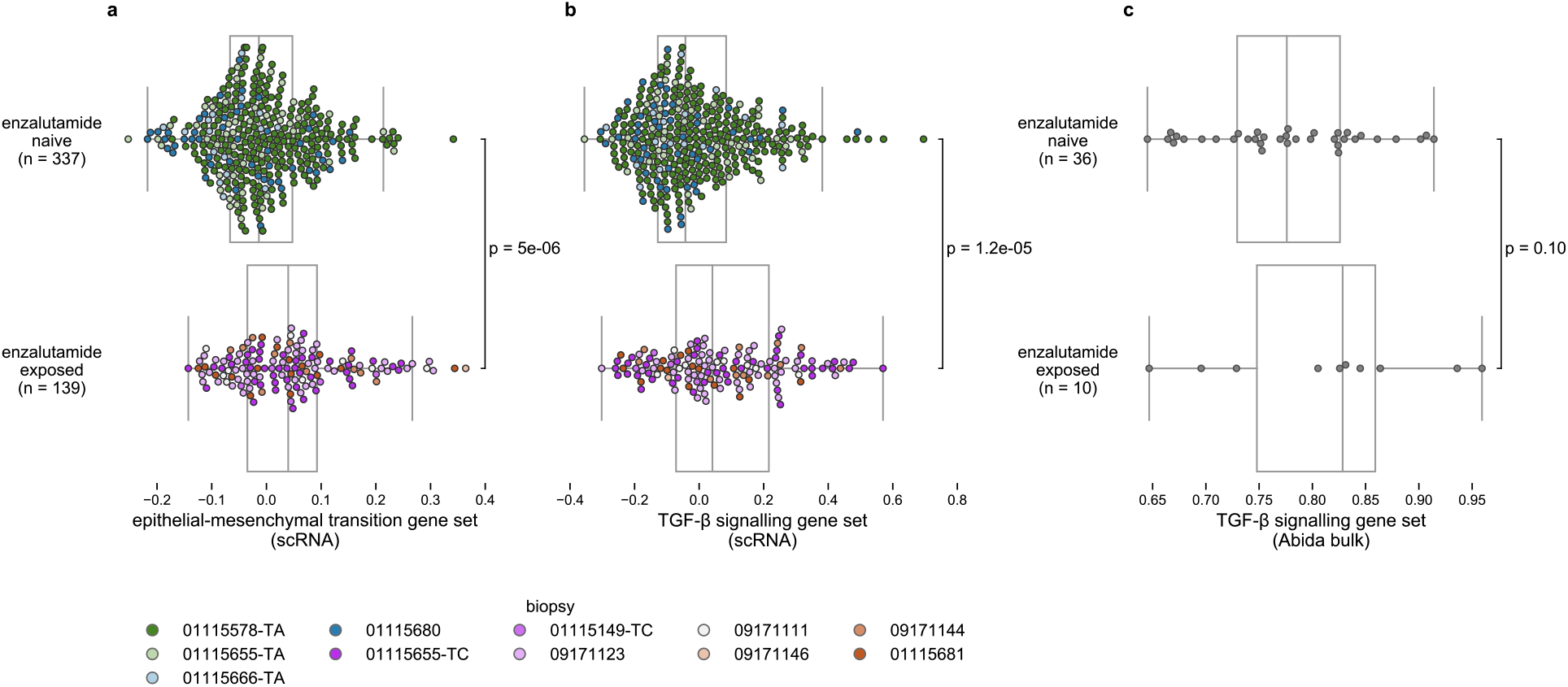
Enzalutamide-exposed adenocarcinoma cells upregulate expression programs associated with epithelial-mesenchymal transition and TGF-β signalling. **a, b)** Hallmark epithelial-mesenchymal transition and TGF-β signalling gene set expression scores for individual cells collected before and after enzalutamide treatment. Each dot represents a single cell and is coloured corresponding to biopsy. *P* values from two-sided Mann-Whitney *U* test. **c)** Hallmark TGF-β signalling gene set expression scores for bulk RNA-seq of prostate adenocarcinoma lymph node biopsies^9^ collected before and after enzalutamide treatment. Each dot represents a single tumour. *P* value from one-sided Mann-Whitney *U* test. Boxplots: centre line: median; box limits: upper and lower quartiles; whiskers extend at most 1.5x interquartile range past upper and lower quartiles.

## Small cell carcinoma regulatory programs

One patient sample within our cohort derived from a small cell carcinoma, a rare aggressive form of prostate cancer that is not responsive to androgen-targeting therapies^23^. As expected, cancer cells from this biopsy differed drastically in their expression programs, with no detectable *AR* expression, strong downregulation of an *AR* regulated gene set, and marked upregulation of a gene set associated with neuroendocrine prostate cancer (Fig. 4a,b; Extended Data Fig. 1)^12,14^.

**Figure 4.**
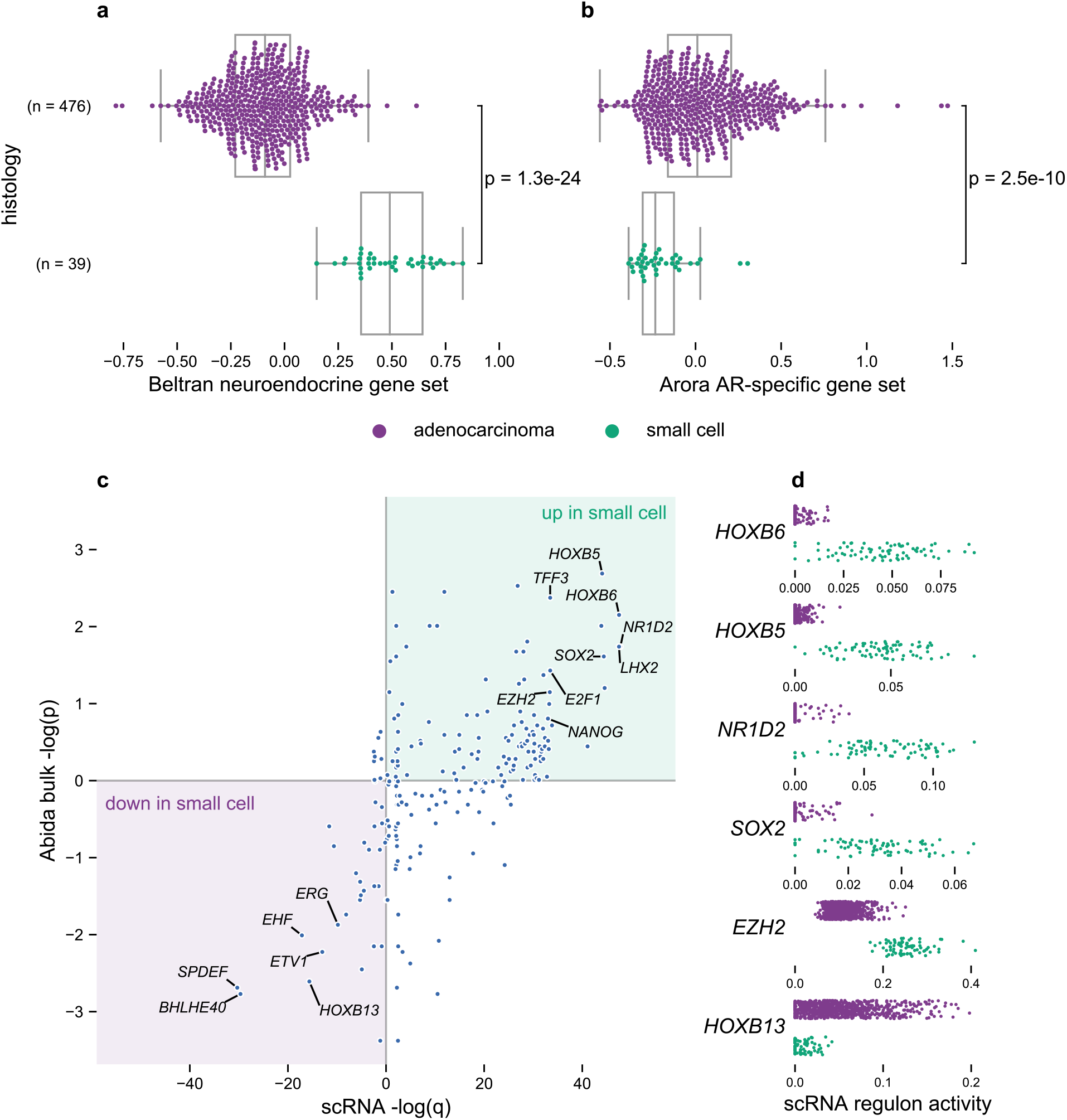
Cancer cells from small cell carcinoma are dominated by distinct regulons compared to adenocarcinoma cells. **a, b)** Gene set expression scores in single cells using an expression signature of neuroendocrine prostate cancer^14^ and of a set of genes under regulation by *AR*^12^. Boxplots: centre line: median; box limits: upper and lower quartiles; whiskers extend at most 1.5x interquartile range past upper and lower quartiles. **c)** Inferred activity of regulons of different transcriptional regulators. x-axis: *q*-values from comparison of inferred regulon activity in cancer cells from small cell carcinoma (n = 76) vs cancer cells from adenocarcinomas (n = 188, sampled as described in Methods) (negative values indicate regulon is less active in small cell carcinoma; two-sided Mann-Whitney *U* test, median outcome of sampling iterations (Methods) with Bonferroni FWER correction). y-axis: *P* values (two-sided Mann-Whitney *U* test, signed as previous) from comparison of expression scores of scRNA-inferred regulons in bulk RNA-seq of small cell carcinomas (n = 8) vs adenocarcinomas (n = 18) from a published cohort^9^. **d)** Regulon activity in single cells for select transcriptional regulators.

To mitigate overestimating the importance of idiosyncratic gene expression patterns from a single biopsy, we inferred transcriptional regulatory factor regulons using all cancer cells from our cohort and compared the inferred regulon activities between small cell carcinoma and adenocarcinoma cells^24^. Additionally, we scored small cell carcinoma and adenocarcinoma bulk transcriptomes from a published cohort for expression of the gene lists inferred to comprise each regulatory factor’s regulon^9,22^ (Methods). Comparing our data and the published cohort, we observed concordant patterns of differential regulon activity between adenocarcinoma and small cell carcinoma (Fig. 4c). Among the transcriptional regulators with decreased activity in small cell carcinoma are *HOXB13*, which mediates *AR* regulatory activity and response to androgens, and *BHLHE40*, previously reported to be regulated by AR^25–27^. Several ETS family transcription factors showed reduced activity in small cell carcinoma, including *ETV1*, which increases prostate adenocarcinoma invasiveness, *EHF*, whose loss confers stem-like features, and *SPDEF*, an AR-regulated transcription factor whose downregulation promotes EMT^28–30^. On the other hand, considering transcriptional regulators with increased regulon expression in small cell carcinoma, we noted the stemness-promoting factors *NANOG* and *SOX2* and the epigenetic regulator *EZH2*, all of which have been reported to promote lineage plasticity and resistance to androgen-targeting therapies^23,31–33^. Among the transcriptional regulators with the most increased activity in small cell carcinoma cells are *E2F1*, which promotes cell cycle progression upon release from RB1 inhibition and is overexpressed in treatment-emergent small cell neuroendocrine prostate cancer and *LHX2*, previously reported in an expression signature of N-myc driven neuroendocrine prostate cancer^34–36^. We also observed increased activity of three transcriptional regulators whose role in small cell carcinoma has not been previously reported: *HOXB5* and *HOXB6*, two homeobox containing transcription factors, and *NR1D2*, a circadian rhythm regulator (Fig. 4c,d)^37^. Thus, even from a single small cell carcinoma case, we recover generalizable patterns of tumour-intrinsic expression differences, implicating both novel regulons and known transcription regulators mediating treatment resistance.

## Cytotoxic cell states and dynamics

To provide a therapeutic axis independent of AR signalling and complementing tumour-intrinsic targeting modalities, clinical trials have tested immune checkpoint inhibitors in prostate cancer. While such therapies have yielded major improvements in a variety of solid tumours, responses in advanced prostate cancer have been muted^1,2^. To improve our understanding of the biology underlying this gap, we characterized infiltrating cytotoxic cells in the mCRPC microenvironment. We sub-clustered T and NK cells identified from initial clustering into 6 clusters, including 2 CD4^+^ T cell populations, 3 largely CD8^+^ T cell populations, and a population of strongly CD16^+^ and largely CD3^−^ cells dominated by NK cells (Fig. 5a; Extended Data Fig. 2a). One population of CD8^+^ T cells chiefly derived from bone biopsies was marked by expression of *CXCR4*, consistent with reports in mice that *CXCR4* is necessary for localization of CD8^+^ T cells to the bone marrow and their subsequent survival^38^ (Fig. 5b; Extended Data Fig. 2a, 3a). This cluster had minimal expression of the effector molecule *GZMB*, while all three other cytotoxic clusters exhibited *GZMB* expression, albeit to varying degrees (Fig. 5b; Extended Data Fig. 2b). Another CD8^+^ T cell population, largely derived from lymph node biopsies, was marked by expression of co-inhibitory receptors *PDCD1*, which encodes PD-1, and *HAVCR2*, which encodes TIM-3, along with elevated expression of *TOX, TIGIT, ICOS, FASLG*, and *LAG3* and minimal *TCF7* expression, suggestive of a dysfunctional effector phenotype (Fig. 5b; Extended Data Fig. 2b,e,f). This population exhibited elevated expression of both *ENTPD1* (encoding CD39, a marker of terminally exhausted CD8^+^ T cells) and *ITGAE* (encoding CD103), whose co-expression identifies infiltrating cytotoxic cells reactive to cancer cells in other human cancers^39,40^ (Extended Data Fig. 2c). Both the NK cell-dominant cluster and the remaining cytotoxic T cell cluster, which included CD8^+^ T cells and likely γδ T cells, were marked by expression of *GNLY* and substantial fractions of cells expressing *CX3CR1* (Fig. 5b; Extended Data Fig. 2d). Cells expressing *CX3CR1* also highly expressed *GZMB* and *PRF1*, consistent with previous reports that *CX3CR1* marks a CD8^+^ T cell population with superior cytolytic function corresponding to a more differentiated effector phenotype that has been observed in models of chronic infection and other cancers^41–43^. We did not observe a distinct cluster of *TCF7* and *SLAMF6* dual-expressing progenitor cells previously reported to mediate response to anti-PD-1 therapy in melanoma (Extended Data Fig. 2e,f)^44^. Broadly, these findings demonstrate that prostate cancer metastases are infiltrated by cytotoxic cells with distinct phenotypes, including dysfunctional and effector states relevant to therapy, that may vary based on metastatic site.

**Figure 5.**
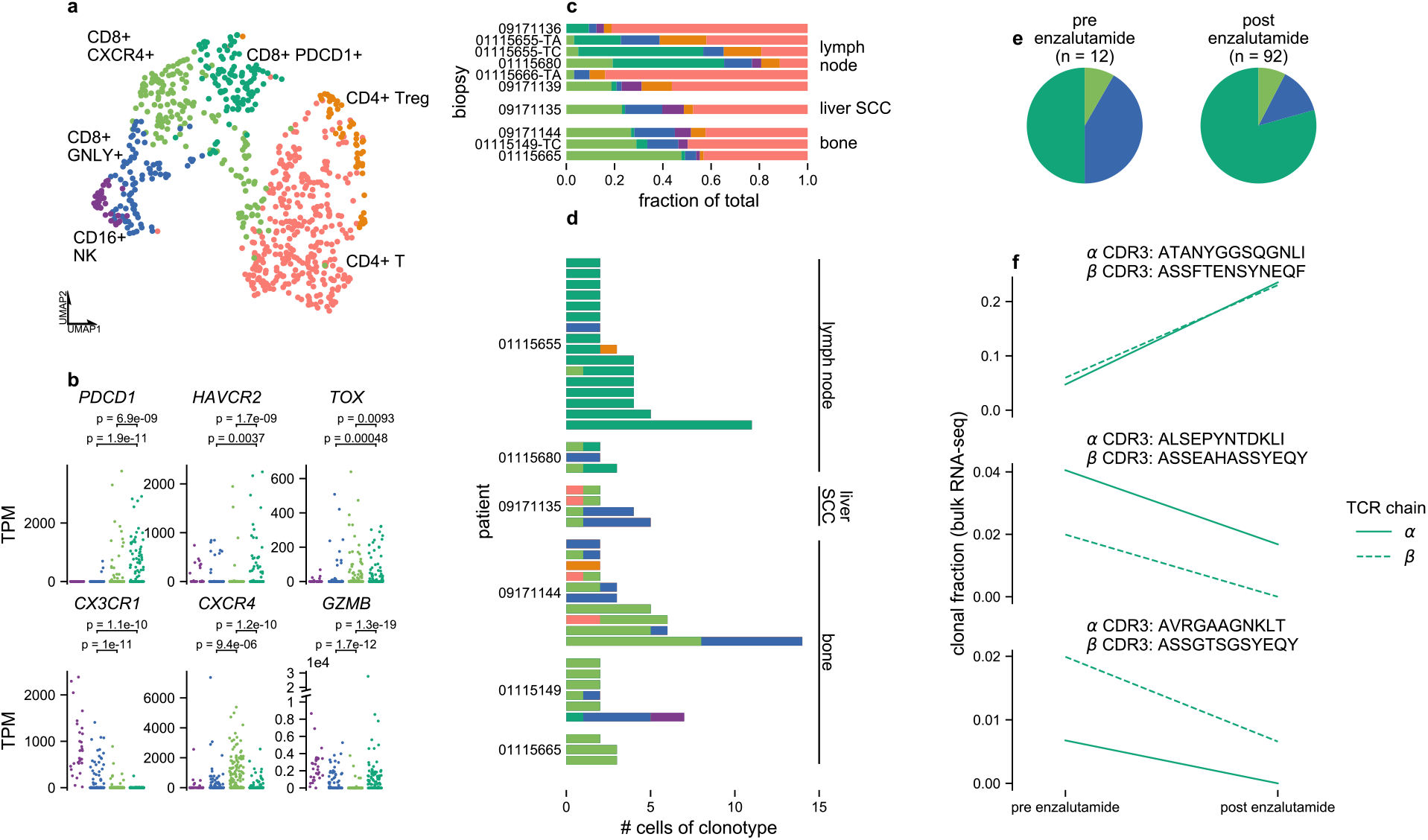
Clonally expanded cytotoxic lymphocytes have different effector phenotypes in distinct metastatic niches. **a)** Sub-clustering of NK and T cells. Each dot represents a single cell projected onto UMAP space coloured corresponding to clustering via the Louvain algorithm. Clusters are manually labelled with dominant phenotype/cell type from patterns of marker gene expression. Cluster colours are used throughout subpanels. **b)** Expression of select marker, effector, and co-inhibitory receptor genes within cytotoxic clusters, CD16^+^ NK (n = 30), *CD8*^+^ *GNLY*^+^ (n = 84), *CD8*^+^ *CXCR4*^+^ (n = 157), and *CD8*^+^ *PDCD1*^+^ (n = 106). *P* values from two-sided Mann Whitney *U* test. **c)** Proportions of cellular phenotypes from each biopsy, grouped by metastatic site, for all biopsies from which high-quality T and NK cells were recovered. **d)** T cell clonotypes from TCR reconstruction. Each bar represents cells sharing a reconstructed productive TCR CDR3 sequence and are grouped by patient. Colours indicate phenotype/cell type. **e)** Proportions of cytotoxic cell phenotypes in patient 01115655 before and after enzalutamide treatment. **f)** Changes in clonal fractions of cytotoxic T cell clonotypes in patient 01115655 following enzalutamide treatment. Each subplot corresponds to a single clonotype with TCRα and β CDR3 amino acid sequences inferred from single-cell RNA-seq. Clonal fractions for the same CDR3 sequences (matching at both nucleotide and amino acid level) inferred from TCR reconstruction in bulk RNA-seq are plotted. All detected single cells of the displayed clonotypes come from the *PDCD1*-expressing CD8^+^ T cell cluster.

Next, we reconstructed T cell receptor (TCR) complementarity-determining region 3 (CDR3) sequences in our scRNA-seq and corresponding bulk RNA-seq data to better understand the clonal dynamics of infiltrating T cells that expand in response to antigen stimulation. Groups of T cells forming part of an expanded clonotype group, indicated by a shared productive CDR3 sequence, were detected in 6 patients. Clonotype groups detected in lymph node metastases were largely comprised of cells from the CD8^+^ T cell cluster with elevated co-inhibitory receptor expression, while clonotype groups detected in bone metastases were largely comprised of cells from the *CXCR4*-expressing CD8^+^ T cell cluster with low *GZMB* expression (Fig. 5d). In one bone biopsy (09171144), a large clonotype group was detected that included both cells from the *CXCR4*-expressing cluster and cells with high *CX3CR1* expression, indicating that cells derived from the same progenitor could take on both phenotypes.

From patient 01115655, we collected cells from biopsies taken both before and after treatment with enzalutamide and noted marked changes in the infiltrating T cell populations (Fig. 5c,e). Before treatment, cytotoxic cells formed a minority of infiltrating T cells, which were dominated by a *SELL*-expressing CD4^+^ T cell population and cells from a CD4^+^ T regulatory cell-enriched cluster with elevated *FOXP3* and *CTLA4* expression (Fig. 5c; Extended Data Fig. 2a). Following treatment, the majority of infiltrating T cells were dysfunctional *PDCD1*-expressing CD8^+^ T cells (Fig. 5c,e). Of note, for the clonotype group with the most cells recovered from this patient, we detected both the corresponding TCRα and TCRβ CDR3 sequences in bulk RNA-seq of biopsies from both timepoints. As inferred from the bulk sequencing data, the clonal fraction increased sharply from ~5% before treatment to ~25% after treatment, making it the largest detected clone (Fig. 5f). All cells of this clonotype group detected in scRNA-seq were part of the *PDCD1*-expressing dysfunctional cluster. Collectively, these observations suggest that CD8^+^ T cells can mount an aggressive response against cancer cells during enzalutamide treatment but also that they take on a dysfunctional phenotype that may limit sustained efficacy.

## Discussion

To overcome limitations in bulk genomic characterization in uncovering cell-type specific contributions to therapeutic resistance in mCRPC, we describe the transcriptomes of individual cells collected from 15 biopsies covering diverse treatment histories, metastatic sites, and histological types. We find that only cancer cell expression significantly explains the sizeable inherited component of prostate cancer risk. Within small cell carcinoma, in addition to recapitulating expression programs promoting lineage plasticity, we identify novel regulators such as *HOXB5, HOBX6*, and *NR1D2*, which show dramatically increased activity both in our study and in an external cohort^9,32,33^. For adenocarcinomas, where resistance to second-generation androgen targeting therapies poses a major clinical challenge, significant attention is devoted to *AR* splice variants encoding constitutively active truncated proteins that promote resistance^7,8^. We find that *AR* splicing varies widely across cells within a single biopsy, with multiple isoforms detectable in individual cells, including those naïve to second-generation androgen targeting therapies. These findings suggest that focused mechanistic understanding of individual isoforms may be insufficient and that additional studies on the overlapping regulatory activity of co-expressed *AR* splice variants are necessary to fully understand their role in therapeutic resistance. More broadly, we identify upregulation of expression programs associated with TGF-β signalling and EMT following exposure to enzalutamide. This is consistent with evidence from pre-clinical models that inhibition of TGF-β signalling promotes reversion of EMT and may sensitize cancer cells to enzalutamide^45,46^. Recent work focused on human mCRPC bone metastases identify tumour associated macrophages as a source of *TGFB1* expression, providing a target cell population for further study and possible therapeutic targeting (Baryawno, N. *et al*. manuscript submitted). Further studies of mCRPC shortly after initiation of enzalutamide may elucidate earlier cellular responses that ultimately precipitate EMT.

Within infiltrating CD8^+^ T cells, a subset expressed dysfunction markers such as *PDCD1*, and this population included cells that underwent a dramatic clonal expansion within a patient after enzalutamide treatment, suggestive of tumour reactivity. The presence of this cell population may explain why some patients with advanced prostate cancer respond to immune checkpoint inhibition in combination with androgen-targeting therapies^47^. *ENTPD1* (CD39) expression in this population suggests that targeting immunosuppressive adenosine signalling may provide benefit in addition to targeting the PD-1 axis^48^. This population was uncommon in bone biopsies, which instead contained clonally expanded CD8^+^ T cells with high effector molecule, low exhaustion marker, and *CX3CR1* expression. This cell state has previously been linked in model systems and other cancers to high cytolytic activity but poor proliferative potential and a requirement for CD4 help^41–43^. Similar cells have been reported as being unresponsive to PD-L1 blockade, potentially explaining the poor performance of immune checkpoint inhibition in mCRPC bone metastases^1,49^. These results highlight the need for additional immunological dissection of mCRPC, where immune checkpoint inhibition has only been indicated for patients with tumour microsatellite instability^1,2^. Importantly, additional investigation should focus on systematic comparisons of bone and lymph node metastases to confirm whether the observed differences in cytotoxic cell infiltration are generalizable. Intriguingly, TGF-β blockade was recently shown to promote response to immune checkpoint inhibition in prostate bone metastases in mice, potentially enabling rational therapeutic combinations to simultaneously act along both androgen and immune axes^50^. Taken together, we report multiple tumour and immune mechanisms across diverse mCRPC metastatic niches that contribute to treatment resistance and provide therapeutic opportunities for this lethal disease.

## METHODS

### Reference versions

We used human genome reference b37 and the GENCODE^51^ release 30 gene annotation lifted over to GRCh37.

### Statistical software

Statistical tests were conducted with SciPy^52^ v1.3.2 running on Python 3.7. R packages were run on R v3.5.1.

### Whole exome analyses

For biopsies with paired tumour and normal samples available, we performed whole exome sequencing with a customized version of a previously described protocol^53^. After DNA shearing, hybridization and exome capture were performed using Illumina’s Rapid Capture Exome Kit (with the exception of the normal sample for 01115149 and the tumour sample for biopsy 01115149-TA, which used the Agilent SureSelect Human All Exon 44Mb v2.0 bait set^54^). Libraries were sequenced with 76 bp paired-end reads on an Illumina instrument.

Reads were aligned using BWA^55^ v0.5.9 and somatic mutations called using a customized version of the Getz Lab CGA WES Characterization pipeline (https://portal.firecloud.org/#methods/getzlab/CGA_WES_Characterization_Pipeline_v0.1_Dec2018/) developed at the Broad Institute. Briefly, we used ContEst^56^ to estimate contamination, MuTect^57^ and Strelka^58^ to call SNVs and indels, DeTiN^59^ to estimate tumour-in-normal contamination, and Orientation Bias Filter^60^ and MAFPoNFilter^61^ to filter sequencing artefacts. Variants were annotated using VEP^62^, Oncotator^63^, and vcf2maf v1.6.17 (https://github.com/mskcc/vcf2maf). Copy number alterations, purity, ploidy, and whole genome doubling status were called using FACETS^64^ v0.5.14. Copy number alterations were evaluated with respect to whole genome doubling status.

### OncoPanel

For biopsies where whole exome sequencing failed, somatic mutation calls, copy number alterations, and ETS fusion status were taken from OncoPanel, a clinical panel sequencing test available at DFCI^65^.

### Sample collection and dissociation for single-cell RNA-seq

Tumour samples were collected and transported in Dulbecco’s Modified Eagle Medium, high glucose, pyruvate (“DMEM”, ThermoFisher Scientific, #11995073) on ice. Single-cell suspensions for single-cell RNA-seq were obtained from tumour core needle biopsies through mechanical and enzymatic dissociation. Samples were first cut into pieces smaller than 1 mm^3^ using a scalpel. For bone biopsies, soft tissue was also scraped from the hard bone surface using a scalpel blade. Samples were then dissociated using one of two protocols, chiefly to optimize for yield of viable cells from different metastatic sites. Cells obtained from the two protocols were comparable, and findings were consistent in sub-analyses of cells processed with the same protocol (Supplementary Fig. 4).

For biopsies, 01115655-TC, 01115666-TA, 01115680, 01115681, 09171111, 09171135, 09171136, and 09171139, the resulting tissue fragments were incubated in 3 mL Accumax (Innovative Cell Technologies, #AM105) for 10 min at room temperature on a rocking shaker (“ACC” protocol). Cell suspensions were then filtered with a 100 μm cell strainer (ThermoFisher Scientific #08-771-19) and spun at 580 g for 5 min at 4°C. In cases where cell pellets appeared bloody, red blood cells were lysed with ACK Lysing Buffer (ThermoFisher Scientific, #A1049201) on ice for 1 min, followed by quenching with PBS and an additional centrifugation. The final cell pellet was resuspended in PBS (Fisher Scientific, #MT21040CV) with 2% FBS (Gemini Bio-Products, #100-106).

For biopsies 01115655-TA, 01115665, 01115149-TC, 01115578-TA, 09171123, 09171144, and 09171146, tissue fragments were incubated in 2-3 mL Medium 199, Earle’s Salts (“M199”, ThermoFisher Scientific, #11150059) with 1 mg/mL Collagenase 4 (Fisher Scientific, #NC9836075), and 10-20 μg/mL DNAse I (StemCell Technologies, #7900) for 5-10 min in a 37°C water bath with intermittent mixing, followed by additional mixing and pipetting (“CD” protocol). Cell suspensions were then filtered with a 100 μm cell strainer, spun at 580 g for 5 min at 4°C, and the resulting pellet resuspended in PBS with 2% FBS. The blood clot from biopsy 09171144 was processed in a similar manner, with the exception that red blood cells were lysed with ACK Lysing Buffer on ice at 5-minute increments for a total of 15 min. For the bone marrow aspirate from biopsy 09171144, mechanical and enzymatic dissociation were not performed, and red blood cells were lysed with ACK Lysing Buffer on ice at 5-minute increments for a total of 10 min.

### Single-cell sorting

Single cell suspensions in PBS with 2% FBS were stained by incubating for 15 minutes at room temperature protected from light with anti-human PTPRC (CD45) monoclonal antibody conjugated to FITC (1:200 dilution, VWR #ABNOMAB12230), anti-human EPCAM antibody conjugated to PE (1:50 dilution, Miltenyi Biotec #130-091-253), and either Calcein-AM (1:200 dilution, ThermoFisher Scientific #C3100MP; biopsies 01115655-TA and 01115665), 7-Aminoactinomycin D (7-AAD) (1:200 dilution, ThermoFisher Scientific #A1310; all other biopsies except sample 01115149-TC), or both (sample 01115149-TC). We first sorted cells with biological dimensions (high FSC-A and high SSC-A), selected single cells, and excluded doublets or triplets (low FSC-W). Next, we sorted live cells (low 7AAD/high Calcein-AM) that were CD45^+^ (high FITC, enriching for immune cells), EPCAM^+^ (high PE, enriching for cancer cells), or double negative (low FITC/low PE, only in biopsy 09171144) (see Supplementary Fig. 5 for example gating). Cell sorting was performed using a BD Biosciences FACSAria cell sorter (IIu or UV) with FACSDiva software. Individual cells were sorted into the wells of 96-well plates with 10 μL TCL buffer (Qiagen, #1070498) with 1% beta-mercaptoethanol (Sigma 63689) per well. Plates were then sealed, vortexed for 10 s, spun at 3,700 rpm for 2 min at 4°C, and frozen on dry ice.

### Transcriptome sequencing, alignment, and quantification

Library preparation for bulk RNA-seq was performed using the Illumina TruSeq Stranded mRNA Sample Preparation Kit (except for biopsy 01115149-TA, which was prepared using the unstranded Illumina TruSeq RNA Sample Preparation protocol (Revision A, 2010)). Libraries were sequenced with 101 bp paired-end reads (except biopsy 01115149-TA, which was sequenced with 76bp paired-end reads) on an Illumina instrument.

For scRNA-seq, RNA was captured from single-cell lysates with 2.2x RNAClean SPRI beads (Beckman Coulter Genomics) without the final elution^67^. After air drying and secondary structure denaturation at 72°C for three minutes, library construction was performed using a slightly customized Smart-seq2 protocol^66^ with 21 cycles of PCR for preamplification. cDNA was purified with 0.8x Ampure SPRI beads (Beckman Coulter Genomics) and eluted in 21 μL TE buffer. During tagmentation and PCR amplification, we used 0.2ng of cDNA per cell and one-eighth of the Illumina NexteraXT (Illumina FC-131-1096) reaction volume. Individual cells were sequenced to a mean depth of ~1.5 million 38 bp paired-end reads on an Illumina NextSeq 500 instrument with 75 cycle high output kits (Illumina TG-160-2005).

After adapter trimming with cutadapt^68^ v2.2, reads were aligned using STAR aligner^69^ v2.7.2b with parameters: --outFilterMultimapNmax 20 --outFilterMismatchNmax 999 --outFilterMismatchNoverReadLmax 0.04 --alignIntronMin 20 --alignMatesGapMax 1250000 --alignIntronMax 1250000 --chimSegmentMin 12 --chimJunctionOverhangMin 12 --alignSJstitchMismatchNmax 5 −1 5 5 --chimMultimapScoreRange 3 --chimScoreJunctionNonGTAG −4 --chimMultimapNmax 20 --chimNonchimScoreDropMin 10 --peOverlapNbasesMin 12 --peOverlapMMp 0.1 --chimOutJunctionFormat 1. sjdbOverhang was set to 1 less than the untrimmed read length. We used multi-sample 2-pass mapping for all samples from each patient, first mapping all samples (bulk and single-cell transcriptomes), merging the SJ.out.tab files, then running the second pass with the jointly called splice junctions. STAR BAMs were passed into Salmon^70^ v0.14.1 to generate gene-level transcript per million (TPM) quantifications with parameters: `--incompatPrior 0.0 --seqBias --gcBias --reduceGCMemory --posBias`. STAR chimeric junctions were supplied to STAR-Fusion^71^ v1.7.0 in kickstart mode to call ETS family fusions.

### Single-cell quality control and clustering

After removing low quality cells (fewer than 500 or more than 10,000 detected genes, fewer than 50,000 reads, or more than 25% expression from mitochondrial genes), we used Seurat^72^ v3.1.0 to perform first-pass clustering using the TPM matrix rescaled to exclude mitochondrial genes. We manually identified and removed a small number of cells with anomalous expression patterns (chiefly co-expression of high levels of haemoglobin with marker genes for non-erythroid cells). Additionally, some cells that did not cluster with erythroid cells (easily identified with dominant haemoglobin expression) nonetheless had low levels of haemoglobin detected, suggestive of contamination from ambient RNA released from lysed erythroid cells. To account for this, we identified genes whose expression was correlated (Pearson correlation > 0.2) with total haemoglobin expression levels in non-erythroid cells with detectable haemoglobin. This consisted of a small set of genes with known function in erythroid cell development and function: *AHSP, GATA1, CA1, EPB42, KLF1, SLC4A1, CA2, GYPA, TFR2, RHAG, FAXDC2, RHD, ALAS2, SPTA1*, and *BLVRB*. To mitigate batch effects driven by different degrees of contaminating ambient erythroid transcripts, we removed these genes, along with the genes encoding haemoglobin subunits, from the expression matrix for all non-erythroid cells.

We repeated the clustering and conducted all downstream analyses with the filtered expression matrix. After joint clustering of all cells (Fig. 1b), we performed sub-clustering on 3 cell subsets: 1) NK and T cells 2) B-lineage cells 3) myeloid cells. We manually labelled clusters by dominant cell identity, as assessed by marker gene expression patterns (Supplementary Table 2). Briefly, cancer cell clusters were identified by expression of *AR*, *KLK3*, or *CHGA;* T cell populations by *CD3D* and *CD3G;* Tregs by *CD4, FOXP3*, and *CTLA4;* NK cells by absence of *CD3D* and *CD3G* and expression of *FCGR3A, FCGR3B*, and *GZMB;* erythroid cells by *HBA* and *HBB*; neutrophils by *ELANE, CEACAM8, AZU1*, and *DEFA1;* macrophages by *APOE, C1QA*, and *C1QB;* monocytes by *ITGAX, CD14, FCGR3A*, and *FCGR3B;* B cells by *CD19* and *MS4A1;* plasmablasts by *CD19* and absence of *MS4A1;* and plasma cells by *SDC1* and high expression of immunoglobulin genes. Additionally, we confirmed the identity of cancer cell clusters by matching transcriptome-inferred copy number alteration profiles generated from inferCNV v0.99.7 (https://github.com/broadinstitute/inferCNV) with those obtained from corresponding bulk whole exome sequencing.

### Cluster specifically expressed genes and LDSC-SEG

We grouped cell clusters into ‘superclusters’ of related cell types (Supplementary Table 2) and performed differential expression to identify markers for each cell cluster, omitting cells in the same supercluster. To mitigate uneven representation of cell types, when comparing against any cluster, we subsampled the same number of cells from each other supercluster and used as even representation as possible of the contained clusters. In determining cancer cell markers, we used as even representation as possible of cells from each biopsy while sampling 200 cancer cells total per iteration. For each cluster, we repeated the sampling 500 times. In each sampling, we performed a one-sided Mann Whitney *U* test for differential expression on all genes with at least 1 TPM expression in at least 10% of the cluster’s cells. We then selected the top 10% most upregulated genes (lowest median *P* value across samplings) as cluster specifically expressed genes. We used a 100kb interval around genes for heritability partitioning with LDSC-SEG v1.0.1, additionally including an annotation corresponding to all genes and the baseline v1.1 model^6^.

### AR *isoform-informative reads*

To identify reads that uniquely map to an *AR* splice variant, we generated a FASTA transcriptome annotation of spliced sequences from isoforms described in literature^7,73–79^. We extracted all reads initially mapped by STAR to the *AR* genomic interval X:66753830-67011796 and then remapped them to our *AR* isoform transcriptome, disallowing clipping, multimapping, or chimeric reads, and requiring end-to-end mapping (STAR parameters: `--outFilterMultimapNmax 1 --alignEndsType EndToEnd --alignSoftClipAtReferenceEnds No --outFilterMismatchNmax 999 --outFilterMismatchNoverReadLmax 0.04 --peOverlapNbasesMin 10`). As our *AR* isoform transcriptome corresponded to transcript sequences *after* splicing, we further excluded reads that mapped with gaps corresponding to additional inferred splice events. We reported all reads that mapped uniquely to an isoform with at most 1 mismatch in Figure 2a.

### Gene set scoring, regulon activity

For both bulk samples and single cells, we scored the activity of gene sets with VISION^22^ v2.0.0. From single cancer cells, we inferred regulons and transcriptional regulatory factor activity with SCENIC^24^ v1.1.2.2. In Figure 4, for single cells, we used SCENIC AUC directly as a measure of regulon activity. For Figure 4c, to infer regulon activity in bulk samples, we extracted the gene sets corresponding to regulons from SCENIC and scored bulk samples for activity of the genes sets using VISION.

When comparing VISION scores in cells from biopsies exposed and naïve to treatment with enzalutamide, we included only cells inferred to be in G1 by Seurat to reduce discovery of signals introduced by different proportions of cycling cells between tumours^72^. We restricted our initial analyses to biopsies with at least 10 G1 cancer cells. As we were interested in generalizable patterns of expression change related to enzalutamide exposure, we attempted to filter out signals driven primarily by expression patterns in any single biopsy by undertaking a subsampling procedure. By considering subsets of the data more balanced for representation from different biopsies, we traded reduced power for more robustness. From either class (enzalutamide naïve vs exposed), we sampled up to 20 cells per biopsy to prevent results from being dominated by tumours with many recovered cells. Additionally, across repeated sampling iterations, we omitted each biopsy in turn, instead sampling cells from other biopsies within its class, keeping the total number of cells the same. We performed 501 iterations of sampling for each biopsy being excluded. For each gene set being scored with VISION, we used the sampling with the median effect size as the summary of all iterations. When measuring effect size, we consistently compared one class vs the other (i.e. always exposed relative to naïve) to ensure consistency in comparisons of direction of effect. We used the corresponding two-sided Mann Whitney *U* test *P* value as the nominal *P* value for the given gene set.

We additionally took the following steps to filter results that appeared to be driven by a single biopsy: for any given biopsy, we compared samplings when cells from the biopsy were held out vs when cells from the biopsy were included. If the proportion of nominally significant results (*P* < 0.05, same direction of effect as the overall median outcome for the given signature) when the biopsy was excluded was less than 80% of the proportion of nominally significant results when the biopsy was included, we considered any overall finding of differential gene set expression as non-robust and did not report it. We reported signatures with FDR <0.05 in Supplementary Table 1^80^. Note that *P* values shown in Figures 3a, 3b, 4a, and 4b are based on all G1 cells and confirmed the findings from this sampling approach.

For comparisons of regulon activity in small cell carcinoma and adenocarcinoma, we took a similar approach, except that in comparing SCENIC AUC scores, we did not restrict to only G1 cells, as the regulons had been inferred with all cancer cells together. As there was one small cell carcinoma biopsy, cells from that biopsy were never selected for omission across samplings.

### Bulk RNA-seq analyses of Abida cohort

In Figures 3c and 4c, we compared our findings to bulk RNA-seq data from a published cohort^9^. Clinical annotations and expression quantifications were obtained from the published supplementary materials and from the authors directly. We converted gene expression values from FKPM to TPM for consistency with the rest of our study. As this cohort included samples sequenced at different centres and from different metastatic sites, we further restricted our analyses to avoid batch effects. For Figure 3c, we analysed only samples sequenced via transcriptome capture at the University of Michigan, as this was the largest identifiably uniformly sequenced subset. For Figure 4c, as the largest number of small cell carcinoma samples were sequenced at Cornell, we included only small cell carcinoma and adenocarcinoma cases from Cornell in our analyses.

### TCR reconstruction

We performed TCR reconstruction and clonotype inference from single-cell RNA-seq with TraCeR^81^ v0.6.0. We performed TCR reconstruction and estimation of clonal fraction from bulk RNA-seq using MiXCR^82^ v3.0.12. TCRs were inferred as detected in both bulk and single-cell RNA-seq if the CDR3 nucleic acid (and therefore amino acid) sequence matched.

## ACKNOWLEDGEMENTS

We thank the patients who participated in this study. This work was supported by the National Science Foundation (GRFP DGE1144152 (M.X.H.)), the National Institutes of Health (T32 GM008313 (M.X.H.), T32 CA009172 (A.K.T), K08 CA234458 (D.L.), K08 CA222663 (B.I.), R01 CA227388 (E.M.V.A), and U01CA233100 (L.F. and E.M.V.A.)), a Burroughs Wellcome Fund Career Award for Medical Scientists (B.I.), the Conquer Cancer Foundation (D.L.), the Society for Immunotherapy of Cancers (D.L.), the Prostate Cancer Foundation (L.F.), a Mark Foundation Emerging Leader Award (E.M.V.A.), a Dana-Farber Medical Oncology Translational grant (A. Rotem and E.M.V.A.), and a PCF-V Foundation Challenge Award (E.M.V.A.). Any opinions, findings, and conclusions or recommendations expressed in this material are those of the authors and do not necessarily reflect the views of the National Science Foundation.

## AUTHOR CONTRIBUTIONS

M.X.H., A. Rotem, A. Regev, M.-E.T., and E.M.V.A. conceived and designed the overall study. M.S.C., A.K., M.-J.S., and C.R. processed the samples and prepared libraries for scRNA-seq. L.D., J.P., A. Rotem, and O.R.-R. supervised and organized the sample collection and preparation. P.S., B.I., and A. Rotem developed the tissue dissociation protocols. Z.Z., L.D., Z.B., and A.K.T. collected the clinical information. M.X.H. and J.C. analysed the bulk exome sequencing data. M.X.H. performed the heritability enrichment analysis. M.X.H. and A.B.-M. analysed *AR* splicing. M.X.H., D.L., and N.I.K. contributed to the analysis of enzalutamide resistance. M.X.H. and K.B. performed the single-cell regulon analyses. M.X.H., K.B., and K.P.B. analysed the immune cells. M.X.H., L.F., M.-E.T., and E.M.V.A. interpreted the data. M.X.H., S.V., and E.M.V.A. wrote the manuscript. All authors reviewed and approved the final manuscript.

---

**Extended Data Figure 1.**
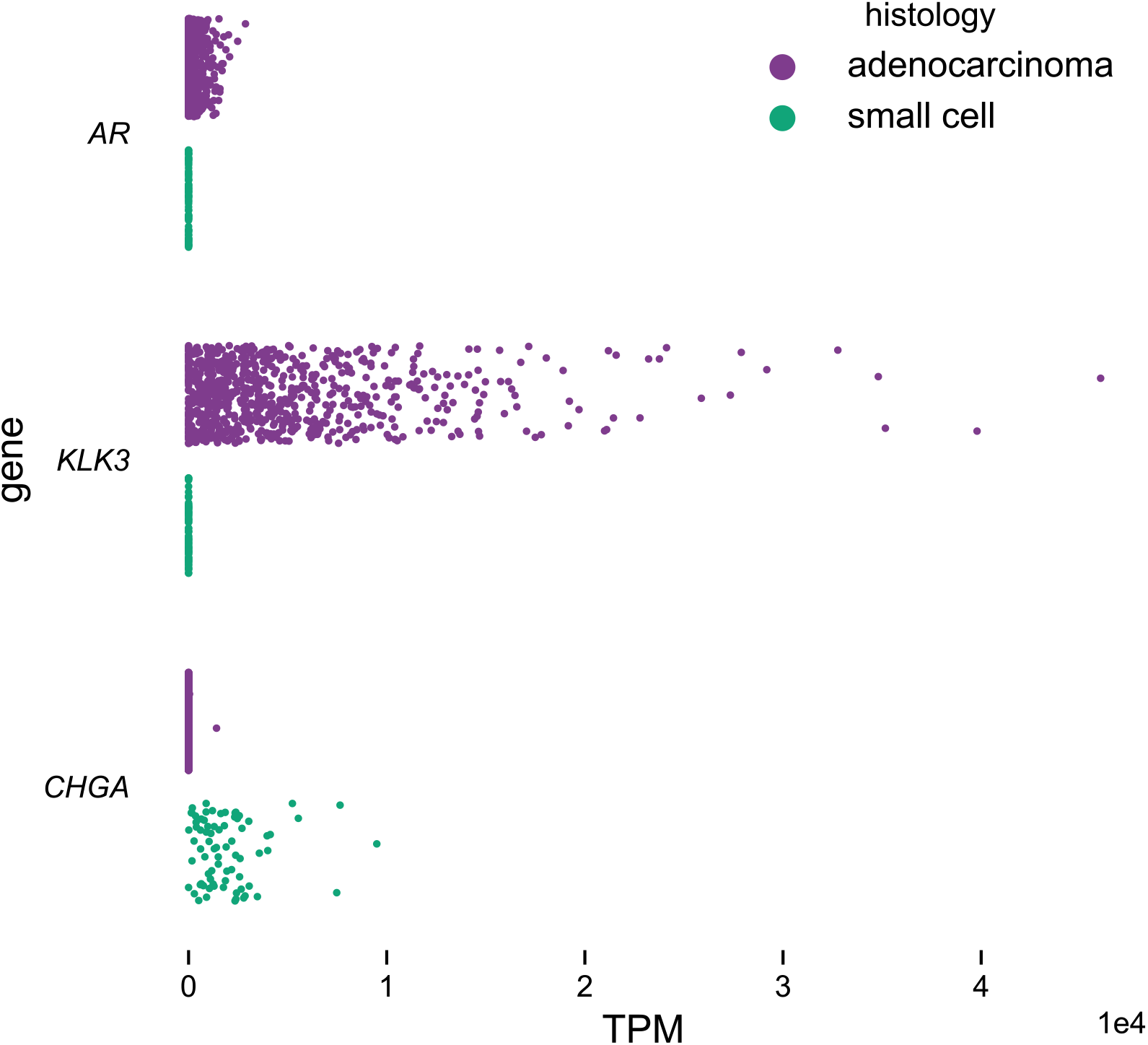
Adenocarcinoma and small cell carcinoma cells are clearly distinguished by marker genes. *AR* and *KLK3* (which encodes PSA) expression marks adenocarcinoma cells (n = 760), while *CHGA* marks small cell carcinoma cells (n = 76).

**Extended Data Figure 2.**
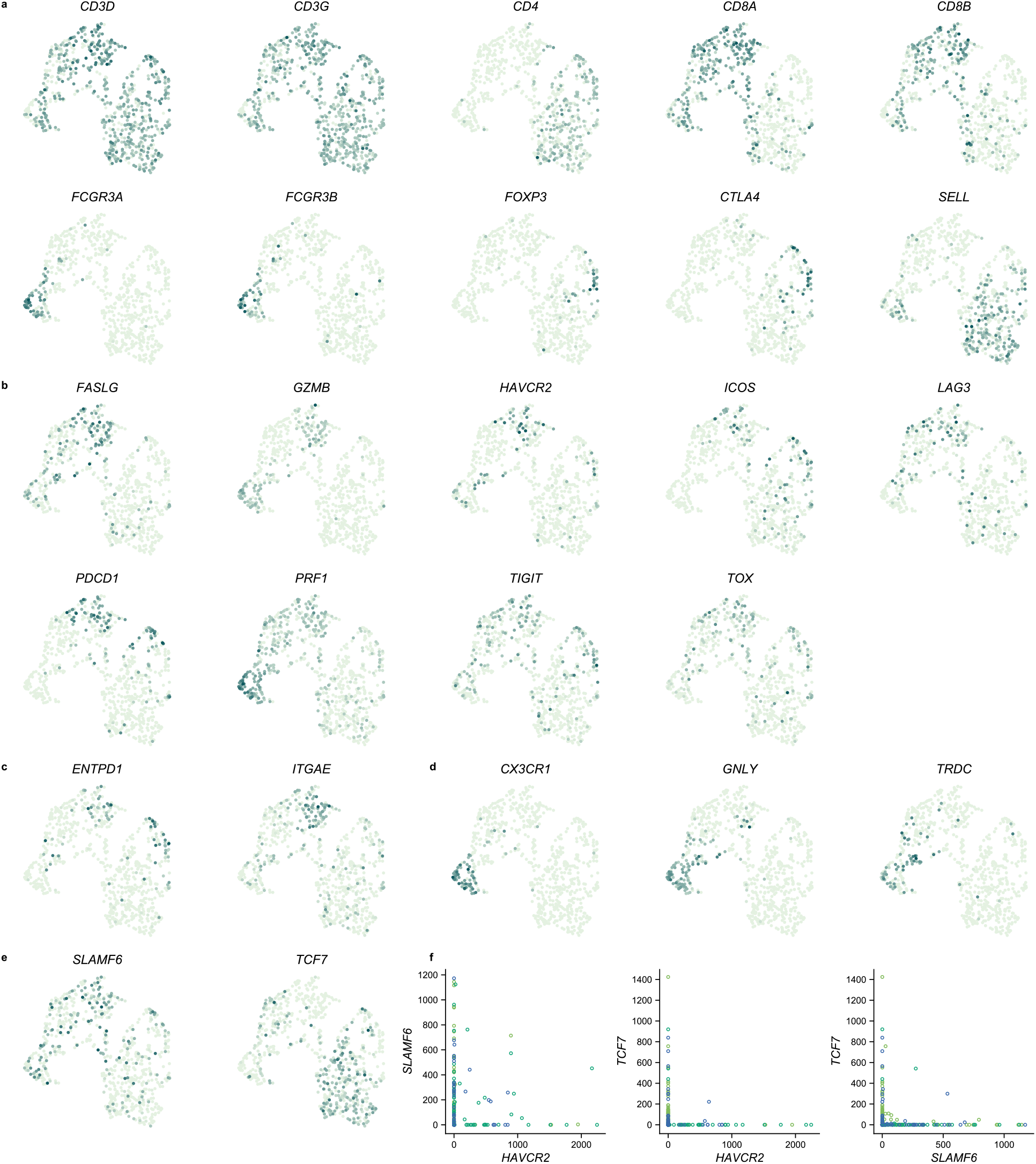
Marker gene expression in NK and T cells. Darker colours indicate higher expression of **a)** cell type markers, **b)** dysfunction and activation markers, **c)** markers of tumour-reactive cytotoxic cells, **d)** genes expressed in a *GNLY*-positive cytotoxic subset, and **e)** genes reported to mark a progenitor population necessary for response after anti-PD-1 therapy in melanoma^44^. Cells are projected onto UMAP space as in Fig. 5a. **f)** Scatterplots showing pairwise co-expression of *HAVCR2, SLAMF6*, and *TCF7* in CD8^+^ T cells. Expression values are in TPM. Points are coloured according to cluster membership as in Fig. 5a.

**Extended Data Figure 3.**
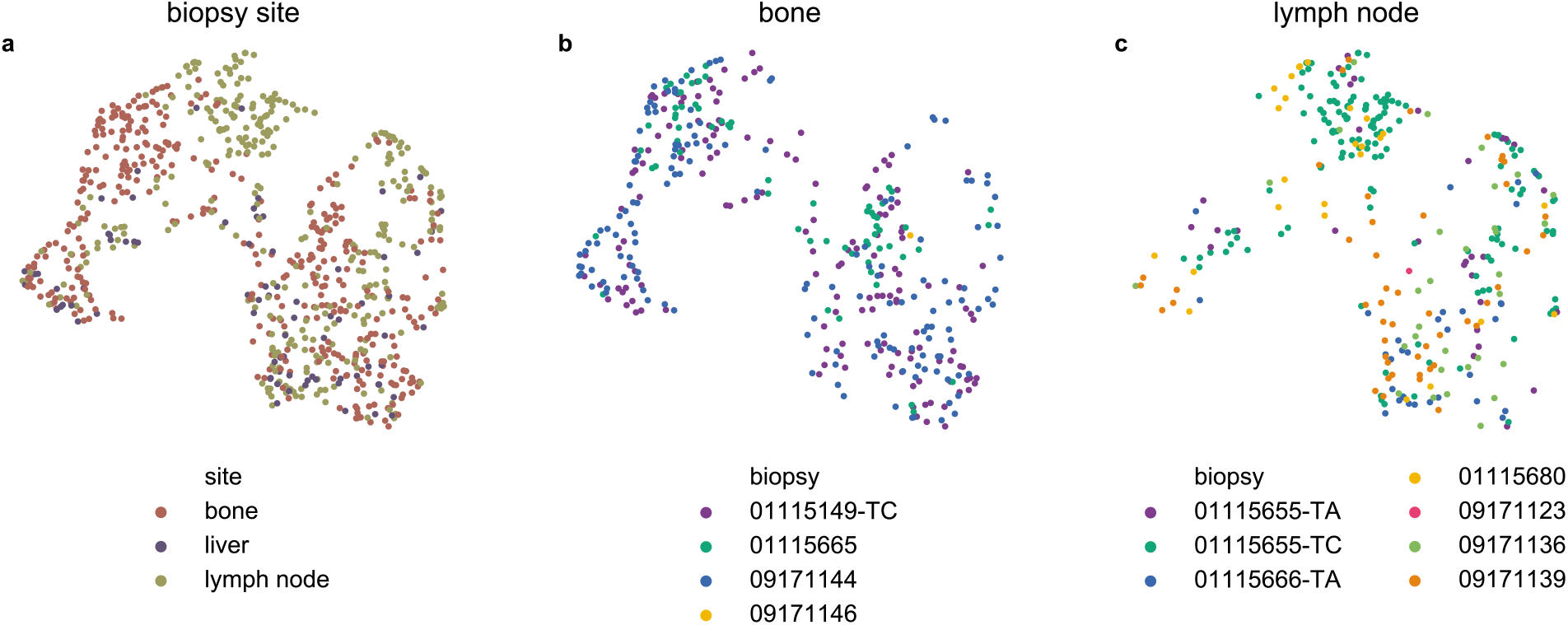
Different cytotoxic subsets are represented in different proportions across metastatic sites. NK and T cells are projected onto UMAP space as in Fig. 5a. **a)** Cells are labelled by site of biopsy. Cells infiltrating **b)** bone and **c)** lymph node metastases are labelled by originating biopsy.

## References

1. Boettcher, A. N. et al. Past, Current, and Future of Immunotherapies for Prostate Cancer. Front. Oncol. 9, (2019).

2. Teo, M. Y., Rathkopf, D. E. & Kantoff, P. Treatment of Advanced Prostate Cancer. Annu. Rev. Med. 70, 479–499 (2019).

3. Picelli, S. et al. Smart-seq2 for sensitive full-length transcriptome profiling in single cells. Nat. Methods 10, 1096 (2013).

4. Mucci, L. A. et al. Familial Risk and Heritability of Cancer Among Twins in Nordic Countries. JAMA 315, 68–76 (2016).

5. Schumacher, F. R. et al. Association analyses of more than 140,000 men identify 63 new prostate cancer susceptibility loci. Nat. Genet. 50, 928 (2018).

6. Finucane, H. K. et al. Heritability enrichment of specifically expressed genes identifies disease-relevant tissues and cell types. Nat. Genet. 50, 621–629 (2018).

7. Lu, J., der Steen, T. V. & Tindall, D. J. Are androgen receptor variants a substitute for the full-length receptor? Nat. Rev. Urol. 12, 137–144 (2015).

8. Antonarakis, E. S. et al. AR-V7 and Resistance to Enzalutamide and Abiraterone in Prostate Cancer. N. Engl. J. Med. 371, 1028–1038 (2014).

9. Abida, W. et al. Genomic correlates of clinical outcome in advanced prostate cancer. Proc. Natl. Acad. Sci. U. S. A. 116, 11428–11436 (2019).

10. Tran, C. et al. Development of a Second-Generation Antiandrogen for Treatment of Advanced Prostate Cancer. Science 324, 787–790 (2009).

11. Liberzon, A. et al. The Molecular Signatures Database Hallmark Gene Set Collection. Cell Syst. 1, 417–425 (2015).

12. Arora, V. K. et al. Glucocorticoid Receptor Confers Resistance to Antiandrogens by Bypassing Androgen Receptor Blockade. Cell 155, 1309–1322 (2013).

13. Hwang, J. H. et al. CREB5 Promotes Resistance to Androgen-Receptor Antagonists and Androgen Deprivation in Prostate Cancer. Cell Rep. 29, 2355–2370.e6 (2019).

14. Beltran, H. et al. Divergent clonal evolution of castration-resistant neuroendocrine prostate cancer. Nat. Med. 22, 298–305 (2016).

15. Li, Y. et al. Targeting cellular heterogeneity with CXCR2 blockade for the treatment of therapy-resistant prostate cancer. Sci. Transl. Med. 11, (2019).

16. Yuan, F. et al. Molecular determinants for enzalutamide-induced transcription in prostate cancer. Nucleic Acids Res. 47, 10104–10114 (2019).

17. Cato, L. et al. ARv7 Represses Tumor-Suppressor Genes in Castration-Resistant Prostate Cancer. Cancer Cell 35, 401–413.e6 (2019).

18. Ragnum, H. B. et al. The tumour hypoxia marker pimonidazole reflects a transcriptional programme associated with aggressive prostate cancer. Br. J. Cancer 112, 382–390 (2015).

19. Hu, R. et al. Distinct Transcriptional Programs Mediated by the Ligand-Dependent Full-Length Androgen Receptor and Its Splice Variants in Castration-Resistant Prostate Cancer. Cancer Res. 72, 3457–3462 (2012).

20. Ertel, A. et al. RB-pathway disruption in breast cancer. Cell Cycle 9, 4153–4163 (2010).

21. Saal, L. H. et al. Poor prognosis in carcinoma is associated with a gene expression signature of aberrant PTEN tumor suppressor pathway activity. Proc. Natl. Acad. Sci. 104, 7564–7569 (2007).

22. DeTomaso, D. et al. Functional interpretation of single cell similarity maps. Nat. Commun. 10, 1–11 (2019).

23. Puca, L., Vlachostergios, P. J. & Beltran, H. Neuroendocrine Differentiation in Prostate Cancer: Emerging Biology, Models, and Therapies. Cold Spring Harb. Perspect. Med. 9, a030593 (2019).

24. Aibar, S. et al. SCENIC: single-cell regulatory network inference and clustering. Nat. Methods 14, 1083–1086 (2017).

25. Navarro, H. I. & Goldstein, A. S. HoxB13 mediates AR-V7 activity in prostate cancer. Proc. Natl. Acad. Sci. 115, 6528–6529 (2018).

26. Norris, J. D. et al. The Homeodomain Protein HOXB13 Regulates the Cellular Response to Androgens. Mol. Cell 36, 405–416 (2009).

27. Kiss, Z. & Ghosh, P. Abstract 2898: Elucidating the role of BHLHE40/DEC1/SHARP2/STRA13 in prostate cancer. Cancer Res. 76, 2898–2898 (2016).

28. Baena, E. et al. ETV1 directs androgen metabolism and confers aggressive prostate cancer in targeted mice and patients. Genes Dev. 27, 683–698 (2013).

29. Albino, D. et al. ESE3/EHF Controls Epithelial Cell Differentiation and Its Loss Leads to Prostate Tumors with Mesenchymal and Stem-like Features. Cancer Res. 72, 2889–2900 (2012).

30. Tsai, Y.-C. et al. Androgen deprivation therapy-induced epithelial-mesenchymal transition of prostate cancer through downregulating SPDEF and activating CCL2. Biochim. Biophys. Acta Mol. Basis Dis. 1864, 1717–1727 (2018).

31. Borges, G. T. et al. Conversion of Prostate Adenocarcinoma to Small Cell Carcinoma-Like by Reprogramming. J. Cell. Physiol. 231, 2040–2047 (2016).

32. Ku, S. Y. et al. Rb1 and Trp53 cooperate to suppress prostate cancer lineage plasticity, metastasis, and antiandrogen resistance. Science 355, 78–83 (2017).

33. Mu, P. et al. SOX2 promotes lineage plasticity and antiandrogen resistance in TP53- and RB1-deficient prostate cancer. Science 355, 84–88 (2017).

34. Lee, J. K. et al. N-Myc Drives Neuroendocrine Prostate Cancer Initiated from Human Prostate Epithelial Cells. Cancer Cell 29, 536–547 (2016).

35. Aggarwal, R. R. et al. Whole-Genome and Transcriptional Analysis of Treatment-Emergent Small-Cell Neuroendocrine Prostate Cancer Demonstrates Intraclass Heterogeneity. Mol. Cancer Res. 17, 1235–1240 (2019).

36. Rickman, D. S. et al. ERG Cooperates with Androgen Receptor in Regulating Trefoil Factor 3 in Prostate Cancer Disease Progression. Neoplasia N. Y. N 12, 1031–1040 (2010).

37. Cho, H. et al. Regulation of Circadian Behavior and Metabolism by Rev-erbα and Rev-erbβ. Nature 485, 123–127 (2012).

38. Goedhart, M. et al. CXCR4, but not CXCR3, drives CD8+ T-cell entry into and migration through the murine bone marrow. Eur. J. Immunol. 49, 576–589 (2019).

39. Duhen, T. et al. Co-expression of CD39 and CD103 identifies tumor-reactive CD8 T cells in human solid tumors. Nat. Commun. 9, 1–13 (2018).

40. Gupta, P. K. et al. CD39 Expression Identifies Terminally Exhausted CD8+ T Cells. PLOS Pathog. 11, e1005177 (2015).

41. Zander, R. et al. CD4+ T Cell Help Is Required for the Formation of a Cytolytic CD8+ T Cell Subset that Protects against Chronic Infection and Cancer. Immunity 51, 1028–1042.e4 (2019).

42. Kanev, K. et al. Proliferation-competent Tcf1 CD8 T cells in dysfunctional populations are CD4 T cell help independent. Proc. Natl. Acad. Sci. 116, 20070–20076 (2019).

43. Hudson, W. H. et al. Proliferating Transitory T Cells with an Effector-like Transcriptional Signature Emerge from PD-1+ Stem-like CD8+ T Cells during Chronic Infection. Immunity 51, 1043–1058.e4 (2019).

44. Miller, B. C. et al. Subsets of exhausted CD8 + T cells differentially mediate tumor control and respond to checkpoint blockade. Nat. Immunol. 20, 326–336 (2019).

45. Paller, C. et al. TGF-β receptor I inhibitor enhances response to enzalutamide in a pre-clinical model of advanced prostate cancer. The Prostate 79, 31–43 (2019).

46. Song, B. et al. Targeting FOXA1-mediated repression of TGF-β signaling suppresses castration-resistant prostate cancer progression. J. Clin. Invest. 129, 569–582.

47. Graff, J. N. et al. Early evidence of anti-PD-1 activity in enzalutamide-resistant prostate cancer. Oncotarget 7, 52810–52817 (2016).

48. Leone, R. D. & Emens, L. A. Targeting adenosine for cancer immunotherapy. J. Immunother. Cancer 6, 57 (2018).

49. Yamauchi, T., Hoki, T., Odunsi, K. & Ito, F. Identification of dysfunctional CD8+ T-cell subsets rescued by PD-L1 blockade in the tumor microenvironment. J. Immunol. 200, 58.3–58.3 (2018).

50. Jiao, S. et al. Differences in Tumor Microenvironment Dictate T Helper Lineage Polarization and Response to Immune Checkpoint Therapy. Cell 179, 1177–1190.e13 (2019).

## Methods References

51. Frankish, A. et al. GENCODE reference annotation for the human and mouse genomes. Nucleic Acids Res. 47, D766–D773 (2019).

52. Virtanen, P. et al. SciPy 1.0: fundamental algorithms for scientific computing in Python. Nat. Methods 1–12 (2020) doi:10.1038/s41592-019-0686-2.

53. Fisher, S. et al. A scalable, fully automated process for construction of sequence-ready human exome targeted capture libraries. Genome Biol. 12, R1 (2011).

54. Gnirke, A. et al. Solution hybrid selection with ultra-long oligonucleotides for massively parallel targeted sequencing. Nat. Biotechnol. 27, 182–189 (2009).

55. Li, H. & Durbin, R. Fast and accurate short read alignment with Burrows–Wheeler transform. Bioinformatics 25, 1754–1760 (2009).

56. Cibulskis, K. et al. ContEst: estimating cross-contamination of human samples in next-generation sequencing data. Bioinforma. Oxf. Engl. 27, 2601–2602 (2011).

57. Cibulskis, K. et al. Sensitive detection of somatic point mutations in impure and heterogeneous cancer samples. Nat. Biotechnol. 31, 213–219 (2013).

58. Saunders, C. T. et al. Strelka: accurate somatic small-variant calling from sequenced tumor-normal sample pairs. Bioinforma. Oxf. Engl. 28, 1811–1817 (2012).

59. Taylor-Weiner, A. et al. DeTiN: overcoming tumor-in-normal contamination. Nat. Methods 15, 531–534 (2018).

60. Costello, M. et al. Discovery and characterization of artifactual mutations in deep coverage targeted capture sequencing data due to oxidative DNA damage during sample preparation. Nucleic Acids Res. 41, e67 (2013).

61. Lawrence, M. S. et al. Discovery and saturation analysis of cancer genes across 21 tumour types. Nature 505, 495–501 (2014).

62. McLaren, W. et al. The Ensembl Variant Effect Predictor. Genome Biol. 17, 122 (2016).

63. Ramos, A. H. et al. Oncotator: cancer variant annotation tool. Hum. Mutat. 36, E2423–2429 (2015).

64. Shen, R. & Seshan, V. E. FACETS: allele-specific copy number and clonal heterogeneity analysis tool for high-throughput DNA sequencing. Nucleic Acids Res. 44, e131–e131 (2016).

65. Garcia, E. P. et al. Validation of OncoPanel: A Targeted Next-Generation Sequencing Assay for the Detection of Somatic Variants in Cancer. Arch. Pathol. Lab. Med. 141, 751–758 (2017).

66. Picelli, S. et al. Full-length RNA-seq from single cells using Smart-seq2. Nat. Protoc. 9, 171–181 (2014).

67. Shalek, A. K. et al. Single-cell transcriptomics reveals bimodality in expression and splicing in immune cells. Nature 498, 236–240 (2013).

68. Martin, M. Cutadapt removes adapter sequences from high-throughput sequencing reads. EMBnet.journal 17, 10–12 (2011).

69. Dobin, A. et al. STAR: ultrafast universal RNA-seq aligner. Bioinformatics 29, 15–21 (2013).

70. Patro, R., Duggal, G., Love, M. I., Irizarry, R. A. & Kingsford, C. Salmon: fast and bias-aware quantification of transcript expression using dual-phase inference. Nat. Methods 14, 417–419 (2017).

71. Haas, B. J. et al. Accuracy assessment of fusion transcript detection via read-mapping and de novo fusion transcript assembly-based methods. Genome Biol. 20, 213 (2019).

72. Butler, A., Hoffman, P., Smibert, P., Papalexi, E. & Satija, R. Integrating single-cell transcriptomic data across different conditions, technologies, and species. Nat. Biotechnol. 36, 411–420 (2018).

73. Robinson, D. et al. Integrative Clinical Genomics of Advanced Prostate Cancer. Cell 161, 1215–1228 (2015).

74. Hu, R. et al. Ligand-independent androgen receptor variants derived from splicing of cryptic exons signify hormone-refractory prostate cancer. Cancer Res. 69, 16–22 (2009).

75. Watson, P. A. et al. Constitutively active androgen receptor splice variants expressed in castration-resistant prostate cancer require full-length androgen receptor. Proc. Natl. Acad. Sci. U. S. A. 107, 16759–16765 (2010).

76. Lu, C. & Luo, J. Decoding the androgen receptor splice variants. Transl. Androl. Urol. 2, 178–186 (2013).

77. Hu, D. G. et al. Identification of androgen receptor splice variant transcripts in breast cancer cell lines and human tissues. Horm. Cancer 5, 61–71 (2014).

78. Hu, R., Isaacs, W. B. & Luo, J. A snapshot of the expression signature of androgen receptor splicing variants and their distinctive transcriptional activities. The Prostate 71, 1656–1667 (2011).

79. Sun, S. et al. Castration resistance in human prostate cancer is conferred by a frequently occurring androgen receptor splice variant. J. Clin. Invest. 120, 2715–2730 (2010).

80. Benjamini, Y. & Hochberg, Y. Controlling the False Discovery Rate: A Practical and Powerful Approach to Multiple Testing. J. R. Stat. Soc. Ser. B Methodol. 57, 289–300 (1995).

81. Stubbington, M. J. T. et al. T cell fate and clonality inference from single-cell transcriptomes. Nat. Methods 13, 329–332 (2016).

82. Bolotin, D. A. et al. MiXCR: software for comprehensive adaptive immunity profiling. Nat. Methods 12, 380–381 (2015).

