## Supplementary Figures 1-5, Supplementary Table 1 for "Transcriptional mediators of treatment resistance in lethal prostate cancer"

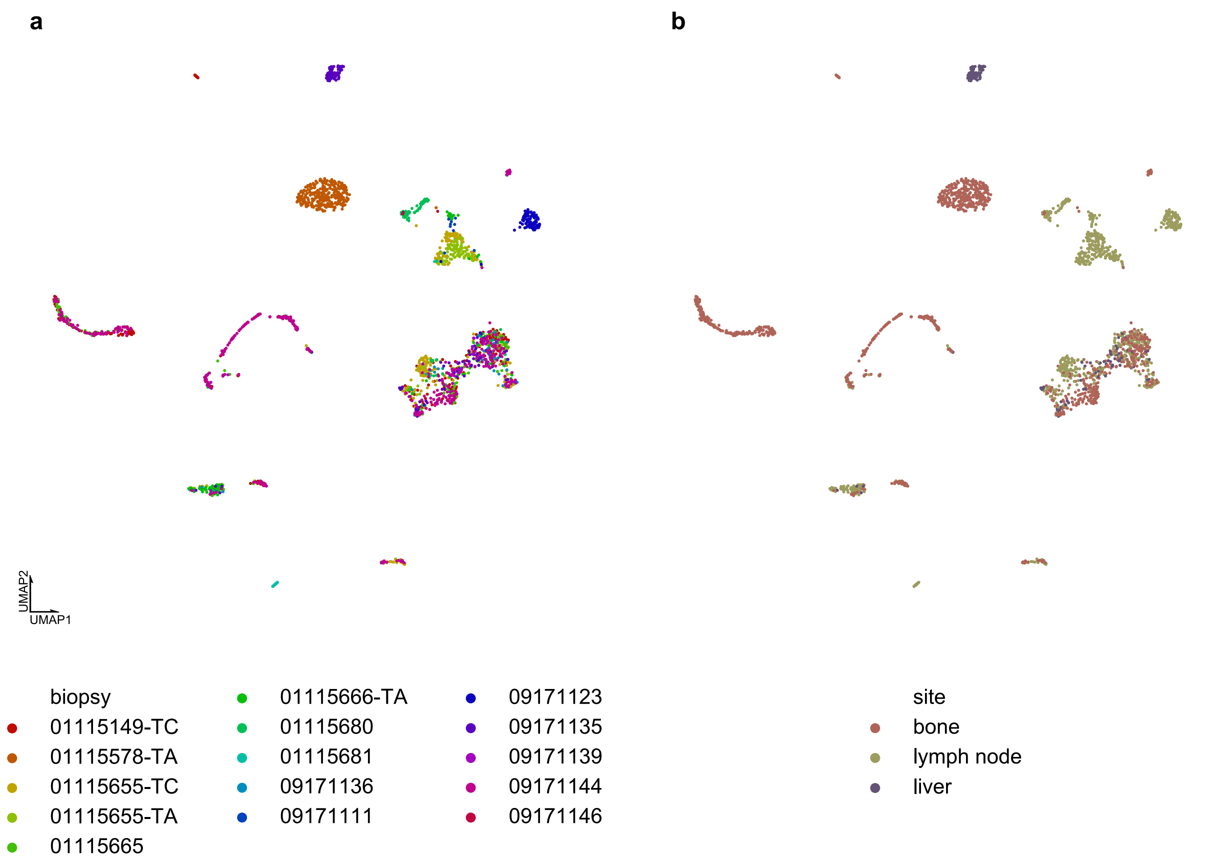


**Supplementary Figure 1.** Projection of single-cell expression onto the first two dimensions of UMAP space.

Cells are coloured corresponding to **a)** biopsy of origin or **b)** metastatic site.

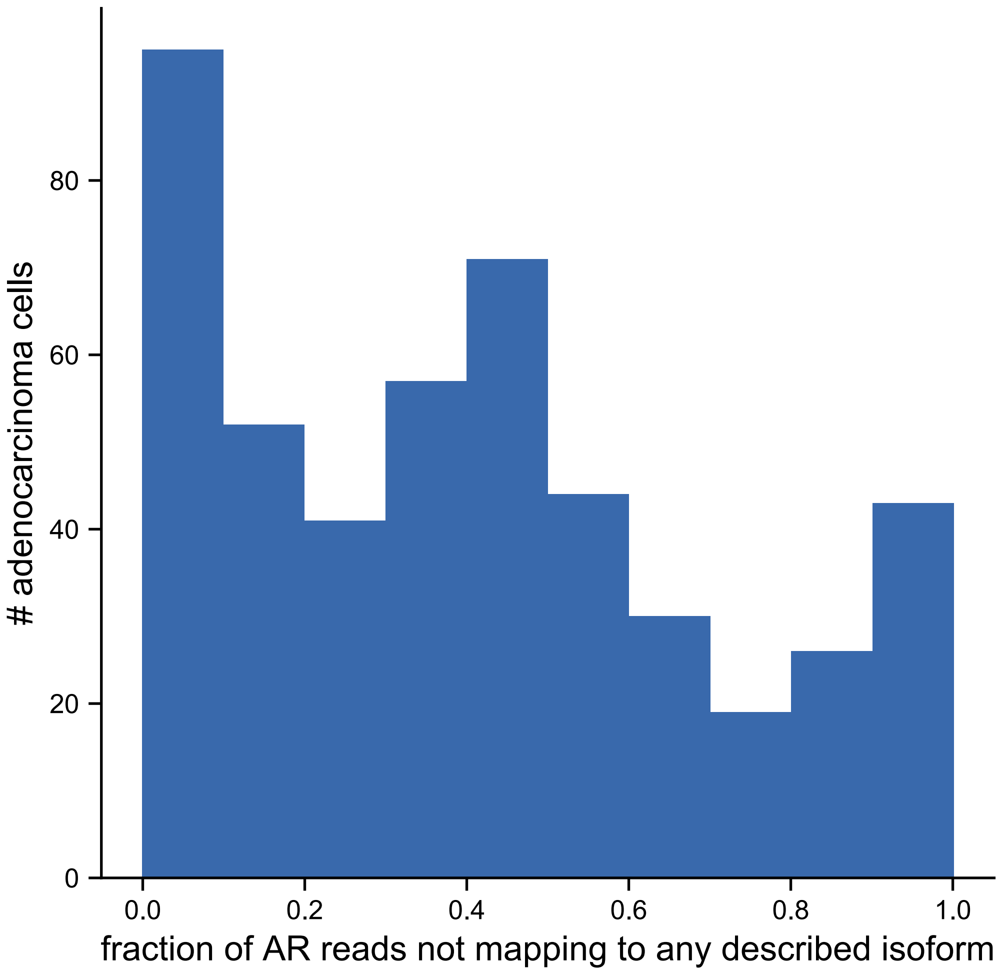


**Supplementary Figure 2.** Significant proportions of reads mapping to the *AR* locus do not map to any literature described *AR* isoform.

| signature | p | exposed vs naive | q |
|---|---|---|---|
| HALLMARK_EPITHELIAL_MESENCHYMAL_TRANSITION | 0.000055 | up | 0.0017 |
| HALLMARK_MYOGENESIS | 0.0011 | up | 0.015 |
| HALLMARK_TGF_BETA_SIGNALING | 0.0013 | up | 0.015 |
| HALLMARK_ESTROGEN_RESPONSE_LATE | 0.0014 | up | 0.015 |
| CREB5[1] | 0.000027 | down | 0.0017 |

**Supplementary Table 1.** Gene sets significantly different between enzalutamide exposed and naïve cells.

Listed gene sets are significant at Benjamini-Hochberg FDR <0.05. Hallmark signatures from MSigDB[2]. *P* values reported are from two-sided Mann Whitney *U* tests from median outcome of comparisons of subsamples of exposed (n = 67) vs naïve cells (n = 76). For details of statistical tests, see Methods.

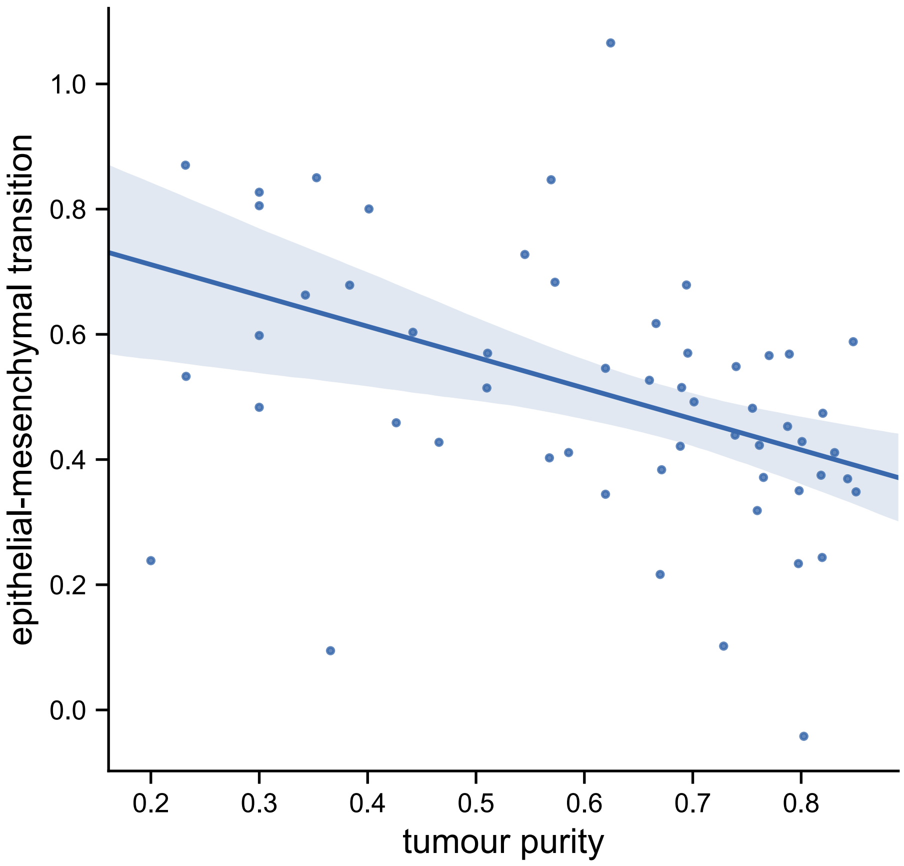


**Supplementary Figure 3.** Tumour purity and hallmark epithelial-mesenchymal signature scores from Abida cohort[3].

Tumour purity was taken from an earlier publication with overlapping samples[4]. All displayed points are from lymph node metastases.

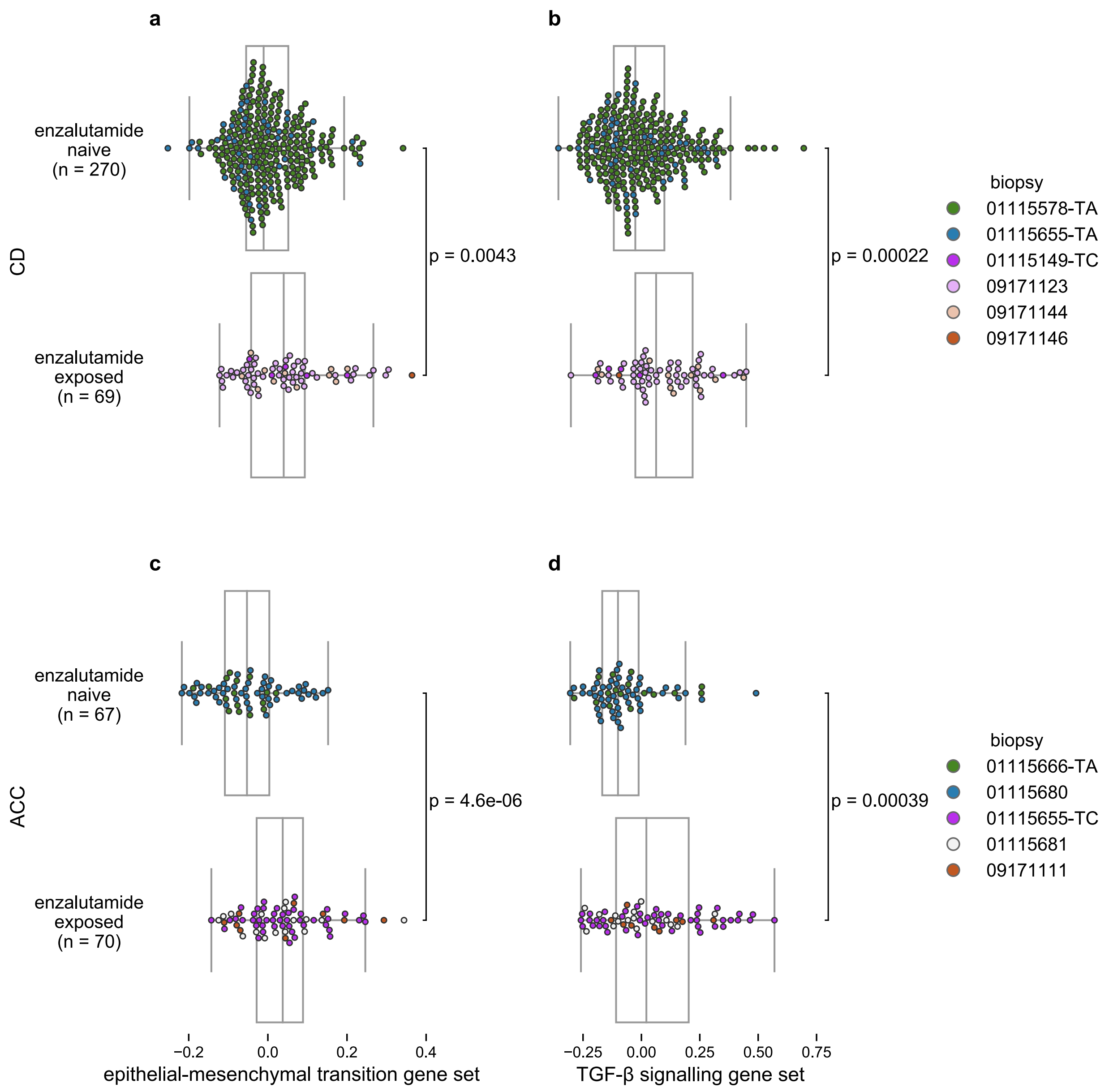


**Supplementary Figure 4.** Changes in expression programs after enzalutamide exposure are concordant across dissociation conditions.

**a, b)** Single-cell VISION[5] signature scores for hallmark epithelial-mesenchymal and TGF-β signalling gene sets for biopsies dissociated with the CD protocol and **c, d)** with the ACC protocol (Methods).

Boxplots: centre line: median; box limits: upper and lower quartiles; whiskers extend at most 1.5x interquartile range past upper and lower quartiles. *P* values from two-sided Mann-Whitney *U* test.

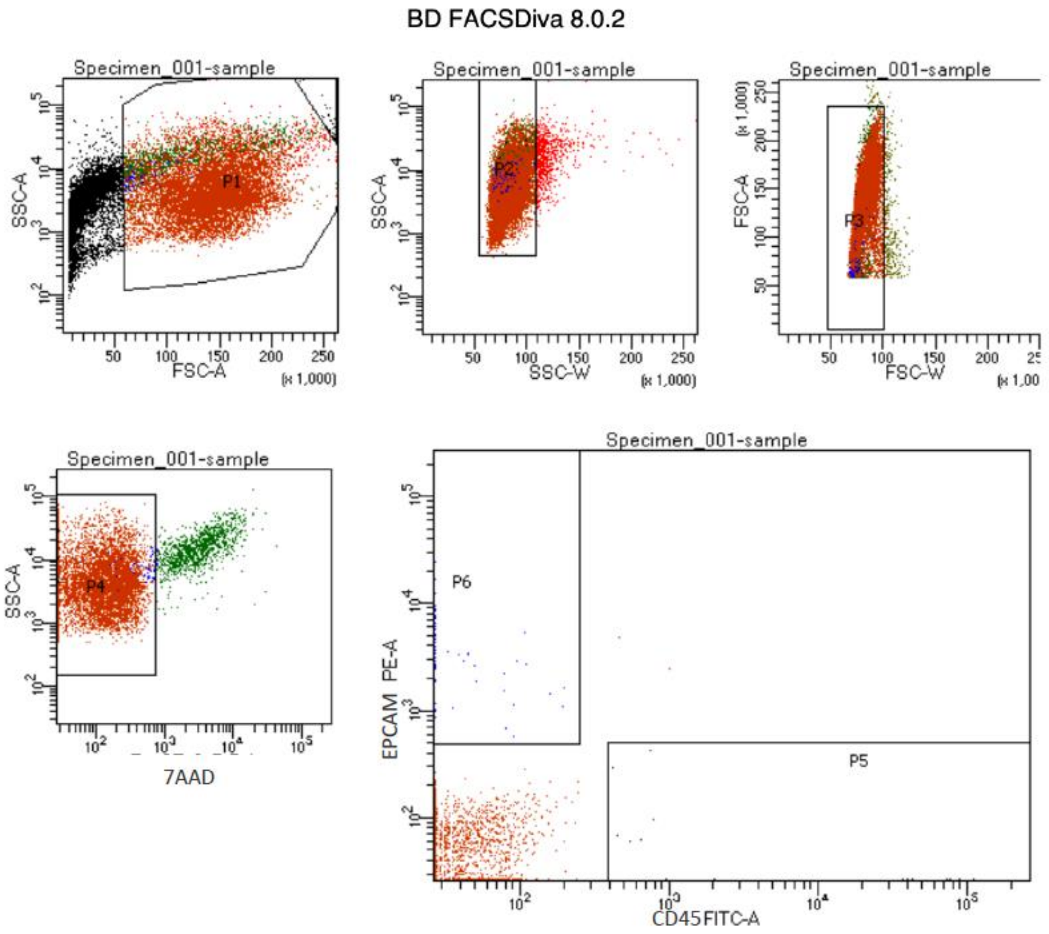


Tube: sample

| Population | #Events | %Parent | %Total |
|---|---|---|---|
| All Events | 20,731 | #### | 100.0 |
| P1 | 11,573 | 55.8 | 55.8 |
| P2 | 10,844 | 93.7 | 52.3 |
| P3 | 10,548 | 97.3 | 50.9 |
| P4 | 9,432 | 89.4 | 45.5 |
| P5 | 15 | 0.2 | 0.1 |
| P6 | 61 | 0.6 | 0.3 |

**Supplementary Figure 5.** Example FACS gating strategy (for biopsy 09171135).

### References


1. Hwang, J. H. *et al.* CREB5 Promotes Resistance to Androgen-Receptor Antagonists and Androgen Deprivation in Prostate Cancer. *Cell Rep.* **29**, 2355-2370.e6 (2019).
2. Liberzon, A. *et al.* The Molecular Signatures Database Hallmark Gene Set Collection. *Cell Syst.* **1**, 417–425 (2015).
3. Abida, W. *et al.* Genomic correlates of clinical outcome in advanced prostate cancer. *Proc. Natl. Acad. Sci. U. S. A.* **116**, 11428–11436 (2019).
4. Rodrigues, D. N. *et al.* Immunogenomic analyses associate immunological alterations with mismatch repair defects in prostate cancer. *J. Clin. Invest.* **128**, 4441–4453 (2018).
5. DeTomaso, D. *et al.* Functional interpretation of single cell similarity maps. *Nat. Commun.* **10**, 1–11 (2019).
